# Dynamic co-existence of bacteriophages and their hosts in the *Arabidopsis thaliana* phyllosphere

**DOI:** 10.64898/2026.04.14.718382

**Authors:** Sheila Roitman, Haim Ashkenazy, Vivienne Hsieh-Wu, Ceren Can, Eva Modly Hurst, Natalie Betz, Katharina Hipp, Detlef Weigel

**Affiliations:** Department of Molecular Biology, Max Planck Institute for Biology Tübingen, 72076 Tübingen, Germany; Institute for Medical Informatics and Bioinformatics, University of Tübingen, 72076 Tübingen, Germany

## Abstract

Bacterial communities and the bacteriophages infecting them are the basis of every ecosystem, including holobionts. The various ways in which these microorganisms interact with each other in complex communities over the life of the host affects the holobiont fitness. Despite being ubiquitous and environmentally relevant, plant-associated microbial communities remain understudied, especially in the phyllosphere, mainly because of the low abundance of microbes and the complexity of the system. In this work we followed bacteria and phage community dynamics in the phyllosphere over a growing cycle of *Arabidopsis thaliana*, to understand the ecology and relevance of bacteriophages in complex bacterial communities. We focused on *Pseudomonas*, a common plant pathogen and commensal, and the phages infecting them, in three setups of increasing complexity: *in vitro,* controlled experiments *in planta* and in wild populations of *A. thaliana*. We found that bacterial communities are resilient to phage infection, and more dynamic than the phages infecting them over the growing season, suggesting that although ubiquitous and abundant, bacteriophages exert selective pressures on leaf bacterial communities only intermittently.

## Introduction

Viruses that infect bacteria, also known as bacteriophages or phages for short, are the most abundant, diverse and ubiquitous entities in the biosphere (Chevallereau et al., 2022; Dion et al., 2020; Puxty & Millard, 2023). Bacteriophages can selectively infect and lyse their bacterial hosts, and upon lysis, release organic matter to be consumed by other members of the microbiome, shifting the balance in microbial populations (Wilhelm & Suttle, 1999). How bacteriophages are actively shaping different ecosystems has been the subject of many studies, first in aquatic environments (Roux et al., 2016; Suttle, 2005), and more recently in soil (Ashelford et al., 2003; Dai et al., 2025; Wu et al., 2025; Xu et al., 2025) and gut microbiomes (Gregory et al., 2020; Guerin et al., 2018; Hsu et al., 2019; Nayfach, Páez-Espino, et al., 2021).

Bacteriophages in the context of plant microbiomes have been mostly studied in the rhizosphere, where bacterial and viral densities are high. Bacteriophages can alter the composition of root-associated microbial communities, thereby alleviating disease symptoms, both by direct predation of the disease-causing pathogens or by un-balancing the soil microbial population (Dai et al., 2025; Wang et al., 2024; Yang et al., 2023; Zhong et al., 2025). Bacteriophages can also reduce disease symptoms when used in seed coating (Erdrich et al., 2024). Additionally, prophages carried by bacteria as integrated mobile genetic elements have been shown to be abundant in plant-associated bacteria (Dougherty et al., 2023; Greenrod et al., 2022; Hulin et al., 2023). Despite these promising findings and the importance of the phyllosphere for primary production, phyllosphere-associated viromes remain poorly characterized. Metagenomics is a powerful technique when dealing with unknown viral diversity, as it allows the identification of phage species and traits present in the viral population. However, characterization of viruses by metagenomics is challenging in the phyllosphere, given the typically low abundance of phages per plant, the high quantities of eukaryotic and microbial DNA, and the accordingly large quantities of starting material needed to create a library. Despite these hurdles, thousands of novel bacteriophages have been lately revealed in the wheat phyllosphere and in the roots of other crops (Dai et al., 2025; Dougherty et al., 2023; Forero-Junco et al., 2022). To understand how many of these metagenomically-assembled genomes and contigs represent active members of the plant microbiome and how important their activity is, it is necessary to isolate and characterize them *in vitro and in planta*.

Microbial communities in the phyllosphere face different challenges than those in the soil, and although they have a fundamentally similar composition (Bai et al., 2015; Bodenhausen et al., 2013), the interplay between microbes and their viruses is bound to be determined by the leaf spatial structure and by abiotic factors. The exterior of leaves in general constitute a harsh environment, with strong UV exposure, poor nutrient availability, and extreme humidity fluctuation, and bacteriophages are only one of the many threats that leaf-associated bacteria must manage. Because of their potential as biocontrol agents, many bacteriophages infecting disease-causing bacteria have been isolated, with phage activity against the bacterial pathogen usually characterized *in vitro*, but sometimes also *in planta* (Erdrich et al., 2025; Feltin et al., 2024, 2025; Korniienko et al., 2022; Rabiey et al., 2020; Skliros et al., 2023; Torres-Barceló et al., 2025). However, it is becoming clear that the success of a phage in rich, well mixed environments, or in gnotobiotic systems does not easily translate to their behavior in realistic, complex microbial communities (Hernandez & Koskella, 2019; Koskella et al., 2022; Svircev et al., 2018). It is thus not surprising that most field trials using phages as biocontrol agents have failed (Svircev et al., 2018). Hence, a more comprehensive understanding of phage-bacteria dynamics in the wild is needed.

Phage activity against plant-associated bacteria has been followed in a handful of natural communities. In cherry trees, for example, it has been shown that added *Pseudomonas*-infecting phages are only present in the leaves as long as their bacterial hosts are, and that the addition of phages does not alter the local microbiota (Papp-Rupar et al., 2025). A study on horse chestnut leaves showed that bacteriophages are more adapted to infect a local bacterial community than phages from other sources (Koskella et al., 2011). There is also a temporal effect, with bacteria becoming resistant over time, while conversely, the infectivity of phages increases (Koskella, 2013, 2014; Koskella & Parr, 2015).

We aim to bridge the laboratory-to-field complexity by simplifying some aspects of the plant-bacteria-phage triangle, while maintaining the complexity of the system as a whole. To this end, we follow bacteria-phage dynamics over a plant’s growth cycle in three settings. First, we infected bacteria with phages *in vitro*, a nutrient-rich environment without spatial structure. Next, we infected bacteria with phages *in planta*, in plants grown in soil under controlled conditions. Finally, we monitor wild plants that harbor a native community of bacteria and phages including lineages closely related to the ones characterized *in vitro*.

As bacterial hosts, we focus on *Pseudomonas*, which are among the most abundant and consistently recovered bacterial species in wild *A. thaliana* phyllospheres in southwest Germany (Karasov et al., 2018; Lundberg et al., 2022; Regalado et al., 2020). *Pseudomonas* are common plant pathogens, but some species are commensals (Bartoli et al., 2014; Shalev, Ashkenazy, et al., 2022). Closely related *Pseudomonas* strains can co-exist in this population over very long time scales, and there is evidence that tailocins (a repurposed phage tail)-mediated antagonistic interactions between strains contribute to this (Backman et al., 2024; Karasov et al., 2018). Conversely, certain commensal strains from this population can protect the plant from pathogenic strains in synthetic community (SynCom) experiments (Shalev, Ashkenazy, et al., 2022; Shalev, Karasov, et al., 2022). In both cases, it seems plausible that bacteriophages play a role in affecting the balance between different *Pseudomonas* strains.

We have isolated and characterized phyllosphere-associated bacteriophages infecting *Pseudomonas* from this population and have monitored bacteria-phage dynamics in SynComs *in vitro* and *in planta*. The phages “followed” their bacterial hosts in the SynCom, affecting the abundance of specific strains, but without a major impact on the overall diversity of the host population. In parallel, we used metagenomics to assess the presence of these and related phages in wild *A. thaliana* phyllospheres, discovering hundreds of novel plant-associated bacteriophages. We followed bacteria and phages during the natural plant growing season. Different from the findings with SynComs, bacteriophages in the wild were more widespread and persistent over the season than their predicted bacterial hosts. Together, our findings suggest that phages are active members of the plant-associated microbiome, yet the extent of their influence might be highly dependent on the exact context, including host physiology and environmental conditions.

## Results

### Bacteriophages of phyllosphere-associated Pseudomonas

The phyllosphere of local *A. thaliana* populations in southwest Germany has been reported to be dominated by *Sphingomonas* and *Pseudomonas* (Karasov et al., 2018, 2024; Lundberg et al., 2022). A large and diverse collection of pathogenic and commensal strains previously assembled (Karasov et al., 2018), and interactions between different members have been characterized in SynCom settings (Shalev, Ashkenazy, et al., 2022; Shalev, Karasov, et al., 2022). Therefore, we decided to focus our phage hunt on these *Pseudomonas* strains, and isolate phages that can be used in a similar SynCom setup. Plants were collected in Tübingen and Gniebl (see Table S1 for coordinates), tissue macerated, resuspended in a buffer and subjected to size selection by ultrafiltration. A dilution-to-extinction approach was followed by three rounds of purification in soft agar plates (see Materials and methods) to obtain pure lysates, which were further characterized by genome sequencing and annotation, imaging by transmission electron microscopy (TEM), and assessment of host range in liquid culture and soft agar.

We isolated and sequenced ten different phages, of which seven were stable over time for characterization (Table S1). All phages but PhC90 have dsDNA and belong to the Caudoviricetes family. PhC90 is a member of the Microviridae with ssDNA. PhC91 belongs to the genus Ventosvirus, PhC60 to the genus Nickievirus, and PhC105 belongs to the Schitoviridae family. Three of the phages, PhC68, PhC94 and PhC95, are jumbo phages, with genomes that are larger than 200 kb and encode the genes characteristic of the Chimalliviridae family (Guan & Bondy-Denomy, 2020). The remaining phages could not be classified. The three jumbo phages, along with PhC91, have a Myovirus morphology, while PhC108, PhC208 and PhC60 have a long non-contractile tail, characteristic of Siphovirus morphology (Figure 1a). Most phages seemed to be purely lytic, although some encode putative integrases and/or recombinases consistent with at least a partial lysogenic lifestyle (Table S2). PhC29 is a lysogenic phage that was activated and enriched in an overnight culture, where the phage entered the lytic cycle, but its particle phase could not be maintained in culture.

**Figure 1.**
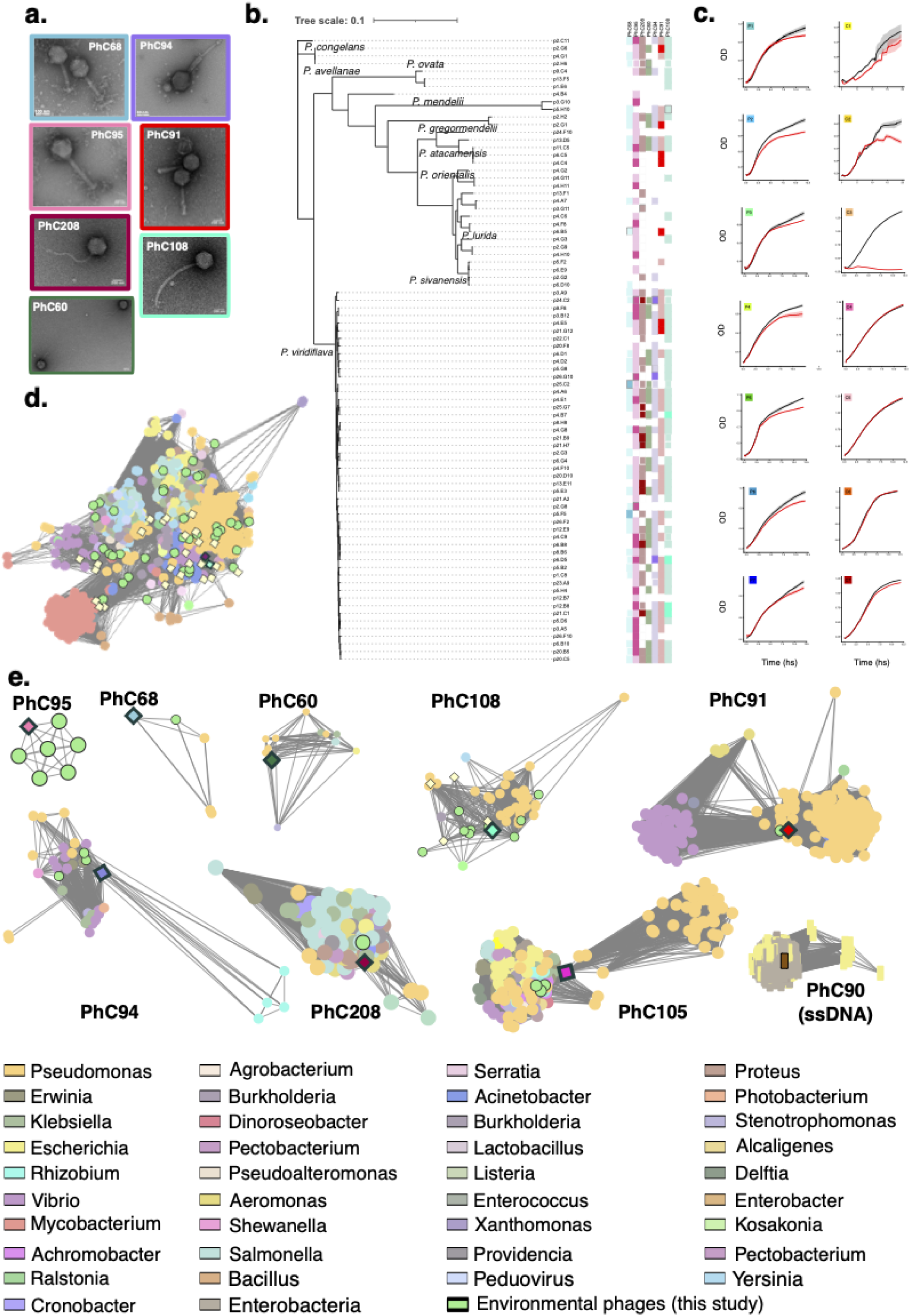
Bacteriophages infecting plant-associated *Pseudomonas*. **a.** Transmission electron microscopy images of newly isolated bacteriophages. For color code of borders, see (b) and (c). Each image has a different scale, as indicated. Full uncropped images can be found in the raw data repository. **b.** Host range of the phages when tested against a representative subset of a local collection of plant-associated *Pseudomonas* from southwest Germany. *Pseudomonas* strains phylogeny based on 127 core proteins. White squares indicate resistance, light color marks delayed or restricted growth, dark color denotes lysis after phage infection. **c.** Phage PhC91 infection curves on individual members of a 14-member *Pseudomonas* SynCom in a liquid assay. OD_600_ was measured every four minutes, cultures were grown in triplicates (SD for each point is given as bars). Black dots are uninfected cultures, red are infected with PhC91 phage at an initial inoculum of 0.05% v/v. Growth curves of the *Pseudomonas* SynCom members infected with all phages can be found in Figure S1. See Table S9 for *Pseudomonas* name equivalences with the phylogenetic tree. **d.** Gene subnetwork of prophages encoded by members of a *Pseudomonas* SynCom (yellow with black contour) and our isolated bacteriophages. Only direct connections to the prophages are shown in this subnetwork. Color code for hosts on the bottom, with orange squares denoting *Pseudomonas*. Shorter edges indicate higher similarity. The two diamonds mark isolate phage PhC108 (aquamarine) and prophage PhC29 (purple). **e.** Gene subnetworks of our isolated phages and known phages. Newly isolated phages are marked with a black contour. A full image of the entire network can be found in Figure S3 and the table used to create it in Table S10.

The host range of bacteriophages can be assessed by several methods. We found that a traditional double-agar layer method for phage infection of *Pseudomonas* (plaque assay) was insufficiently reproducible for our system, as we obtained different outcomes depending of parameters such as ambient temperature, agar percentage, experimenter and bacterial culture used: in some instances, two separate overnight cultures inoculated from the same colony could be either susceptible or resistant to the same phage. Therefore, we complemented plaque-assays with an alternative method for assessing host sensitivity, using OD measurements of liquid cultures, which is gaining popularity due the ability to scale it and being quantitative.

We found *Pseudomonas* strains to have diverse degrees of sensitivity to the phages (Figures 1b, S1). All phages analysed have a broad host range, infecting and/or inhibiting growth of several *Pseudomonas* species and strains (Figure 1b). This is in accordance with new evidence of bacteriophages having a broader host range in the wild, even when interactions could not be easily detected in the laboratory (Bignaud et al., 2025). Most of the infected bacterial strains were only delayed in their growth by the addition of phages, reaching a lower OD at stationary phase, or taking longer to reach it, while some the hosts were completely lysed by at least one phage (Figures 1c, S1).

Because we were interested in measuring the effect of phages in an established SynCom composed of fourteen strains (Shalev, Karasov, et al., 2022) we also characterized their prophage content, as lysogenic phages are integral members of viromes (Dougherty et al., 2023; Knowles et al., 2016; Weitz et al., 2019). Eight strains encode for at least one prophage and eleven encode a tailocin (Table S3, Figure S2). To better characterize the virome in our planned SynCom we used a network analysis to compare their gene content to those of all known Caudoviricetes phages. The SynCom prophages clustered in a large, central node, mostly along phages infecting *Pseudomonas* and other common bacterial species (Figure 1d). This clustering might be the result of these prophages having relatively small genomes (30 - 64 kb), with only a few accessory genes, suggesting a common “core” set of phage metabolic and structural genes. In contrast, for the lytic phages in our SynCom several connectivity patterns can be distinguished (Figure 1e). Within the jumbo phages, PhC95 clustered solely with phyllosphere-metagenomic contigs sequenced in this study, PhC68 clustered with a metagenomic contig and two other *Pseudomonas*-jumbo phages, while PhC94 forms a hub connecting diverse phages infecting several bacterial species. These findings suggest that our jumbo phages have distinct infecting strategies, consistent with their different host ranges (Figure 1c). PhC95 successfully infects several *Pseudomonas* strains, lysing many of them, while PhC68 and PhC94 have a more restricted host range, and are overall less virulent. The low connectivity between these phages in the gene network suggests that the pangenome of jumbo phages is still largely unknown. Our other dsDNA isolates, which cluster mostly with known *Pseudomonas*-infecting phages, are directly connected to phages infecting several common bacterial species in the plant microbiome (Figure 1e). A separate network was constructed for ssDNA phages, PhC90 clusters with other Microviridae phages, an expected outcome given the conserved gene content of this family.

### Relationship of laboratory isolates to phages in phyllosphere metagenomes

A preliminary analysis of the newly isolated phages indicated limited similarity to other phages, especially for the large jumbo phages, and most could not be assigned to a known family. We therefore complemented our isolation efforts with metagenomics of the same *A. thaliana* populations from which the phages had been isolated. We sequenced DNA extracts of leaf samples collected during the 2024-2025 season and assembled reads from individual sampling dates and sites, but pooled across individuals, *de novo*. All identified viral contigs were annotated and checked for completeness, and contigs classified as “complete” and “high quality” were selected for further analyses, including host prediction, taxonomic annotation and clustering with other phage genomes based on gene content and similarity.

Two phages from our collection PhC108 and PhC60 were detected in the metagenomes, as well as phages closely related to PhC95. The gene content and similarity network shows that several of our metagenomically-assembled phyllosphere-associated phages (bright green, black contour) are directly connected to phages infecting various bacterial hosts from the gut of humans and other animals, soil and marine environments (Figure S3). Many of our metagenomically-assembled phages clustered within each other or stayed singletons, showing that the diversity of phyllosphere-associated phages is far from complete. To better characterize these phages we tried to assign them to a known taxonomy, yet out of the manually curated ∼300 high-quality and complete contigs in our metagenomes, only four could be assigned to a genus by the TaxMyPhage algorithm (Table S4) and fourteen to a family by geNomad (Table S5). Some of the other contigs (medium and low quality) could be assigned a taxonomy and a specific host, common bacteria in plant microbiomes (Tables S6, S7). These phages encode multiple common auxiliary metabolic genes (AMGs), such as genes encoding heat shock proteins, MazG, Nrd-proteins (A,B,D,I), a putative phosphate transporter (PhoH), sporulation proteins (P,D,B), or thioredoxins, etc, and some rarer AMGs, such as genes for subunits of cytochrome cd1, a biofilm development protein, copper-related enzymes, CspA, Fumarase D, metallo-dependent phosphatases, OMPA-like proteins, TolA and TolB, TelA (Goldsmith et al., 2011; Rihtman et al., 2019; Schwartz et al., 2022; Sharon et al., 2011; Tahan et al., 2025). These phages also encode several defense systems against bacteriophages, pathogenicity agents such as toxin-antitoxin systems (for example, MazE, YdaS, BrnA, DarT1-associated NADAR, hicA/hicB, ParE-like, Rel-B, Zeta, vbhA), bacteriocins, LemA proteins, pemK, and more (Table S8).

### Bacteriophage proliferation in a SynCom setting

We had tested the host range of our phages on bacterial isolates that came from the same local *A. thaliana* population in southwest Germany as the phages. This in turn afforded us an opportunity to test the effects of these phages on a biologically relevant community of bacteria. With a *Pseudomonas* SynCom that had been previously designed to investigate bacteria-bacteria interactions (Shalev, Ashkenazy, et al., 2022; Shalev, Karasov, et al., 2022), we carried out a time course experiment to understand the temporal dynamics of interactions between phages and their bacterial hosts (Figure 2a). The SynCom included seven closely related pathogenic *P. viridiflava* strains from the ATUE5 clade and seven diverse commensal *Pseudomonas*, with the commensals being able to partially protect the host from the effects of the pathogens (Shalev, Karasov, et al., 2022). These strains had been previously genetically barcoded, so that the abundance of each strain could be easily followed (Shalev, Karasov, et al., 2022) (see Table S9 for name equivalences). We used five treatments, an uninfected control (C), a pathogenic community (seven ATUE5 strains – PC), a mixed community (all 14 strains – MC), the pathogenic community plus lytic phages (PP) and the mixed community with lytic phages (MP). Three weeks after germination, plants from two *A. thaliana* (*Col-0* and *Schl-7*) were inoculated with an airbrush, and grown for another three weeks. Every few days, we sampled four individual plants for each treatment, for a total of 36-48 individuals per treatment (see Table S11 for details). Bacterial and phage abundance was measured by DNA extraction followed by amplicon sequencing and quantitative PCR. In addition, we extracted RNA to follow the expression of 16S rRNA of bacteria and of phage marker genes.

**Figure 2.**
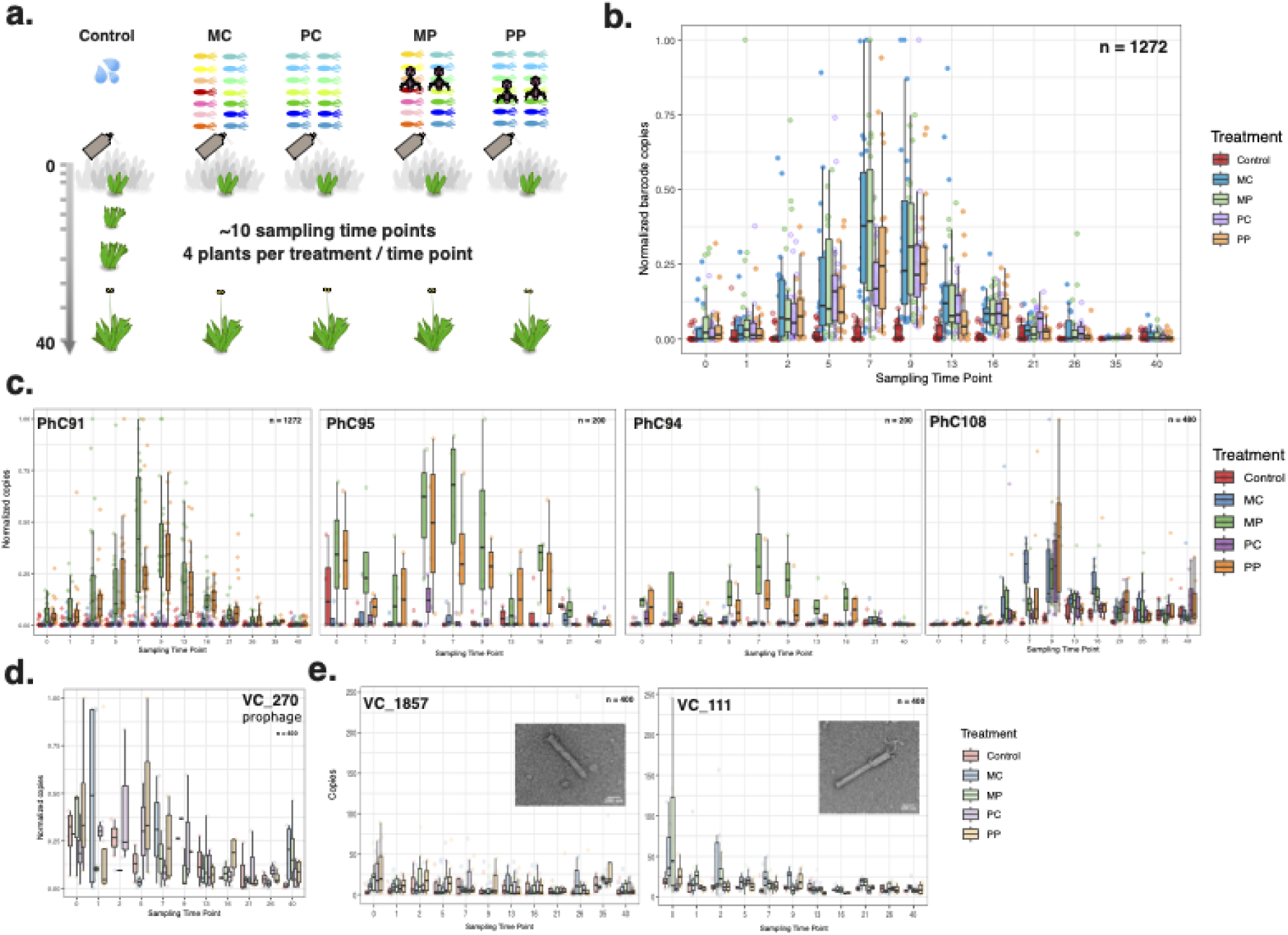
Abundance of bacteria and bacteriophages in soil-grown plants under controlled conditions. **a.** Experimental scheme. Five treatments of barcoded plant-associated *Pseudomonas sp.* communities were prepared: Control (10 mM MgSO_4_), MC – mixed community, PC – pathogenic community, MP – mixed community with added bacteriophages, PP – pathogenic community with added bacteriophages. The five communities were sprayed onto 21-22 day-old *A. thaliana* plants with an airbrush. Four plants per treatment were sampled at each of ten time points over forty days after infection. Sampled plants were weighed, flash frozen and stored until DNA and RNA extraction. **b.** Total abundance of SynCom *Pseudomonas* strains over time, as measured by qPCR on the shared barcode sequence. Each dot represents a plant, colored by treatment. n = 1,272. **c.** Abundance of lytic bacteriophages PhC91, PhC95, PhC94 and PhC108 over time. The number of plants is given on the top right in each subpanel. See Table S12 for absolute abundances. **d.** Expression levels of the prophage VC_270 of *P. viridflava* p4.C9 (P1). n = 400. **e.** Expression levels of one of the tailocin genes over time. n = 400. TEM images are from SynComs grown in vitro. Full uncropped images can be found in the raw data repository.

After having peaked seven to nine days after infection, bacteria decreased, often close to extinction, regardless of treatment and experiment (n = 1,272 plants, from six replicate experiments) (Figure 2b). This was also true for the uninfected controls that are colonized solely by bacteria found in the soil or air (Figure S4), indicating that the demise of the SynCom members is not due to the lytic phages or unique to our focal strains, but instead reflects some feature of our laboratory setting, as this is not the case in wild plants that we examined (see discussion below). Other studies, including one that monitored spontaneous phyllosphere colonization over time in the greenhouse (Maignien et al., 2014), generally reported an increase in bacterial load over the growth cycle. Potentially relevant is that *A. thaliana* plants grown in short days transition from the juvenile to the adult stage at about two weeks of age, and the vegetative-to-reproductive transition starts at about four weeks of age (seven days post infection); this transition can drive microbiome remodelling in roots (Edwards et al., 2018). Measurement of 16S rRNA abundance showed that the plants infected with *Pseudomonas* had orders of magnitude more active bacteria than the control until day 21 post-infection, confirming that our SynCom strains are the dominant and active members of the plant phyllosphere during the relevant span of our experiments (Figure S5). Finally, we note that despite using equal or higher bacterial densities in our setup compared to previous experiments in our laboratory (Shalev, Ashkenazy, et al., 2022; Shalev, Karasov, et al., 2022), the pathogenic community did not cause obvious symptoms including reduced growth of the plants, with their average weight remaining similar for all treatments at each time point (Figure S6).

Among the phages, PhC91 was able to proliferate the most, succeeding to establish and reproduce in all six replicates (Figure 2c). Its levels followed the total abundance of *Pseudomonas*, peaking at days 7-9 and decreasing afterwards down to almost extinction at day 40 post-infection. Two out of three of the jumbo phages added in the phage cocktail managed to reproduce and become established in only one of the replicate experiments, to lower abundance than PhC91 (Figure 2c, Table S12). PhC108, a small siphophage, also managed to establish and replicate in the two replicate experiments where it was added (Figure 2c).

We were surprised to detect PhC108 not only when we added it as part of our phage cocktail (PP and MP), but also in the other two *Pseudomonas* treatments (PC and MC). However, it was not detectable in the uninfected control. We infer that PhC108 must be present in the environment, almost certainly the soil used for the experiments, either as an integrated phage in bacterial cells or as a free viral particle, and once we added *Pseudomonas* communities that included known PhC108 hosts, they could proliferate *in planta*. How exactly PhC108 manages to be transmitted from the apparent soil reservoir to the phyllosphere, i.e., directly or via bacteria, is unknown. In the gene-similarity networks, PhC108 is the only of our lytic phages that clustered together with the prophages in our bacterial collection, suggesting it might have a lysogenic stage in its life cycle, which in turn could explain its reaching the phyllosphere as a hitchhiker in bacterial genomes.

The similar growth pattern of bacteria and phages suggests that these lytic phages are not likely to drive the composition of the leaf *Pseudomonas* communities at the plant level, but are rather “followers”, responding to the changes in the microbial population, as recently suggested for complex environments (Castledine & Buckling, 2024; Papp-Rupar et al., 2025).

Seven members of our SynCom host prophages in their genomes (Table S3, Figure S2), but we could rarely detect their activation in a conclusive manner. One of the few examples of an activated prophage is the viral cluster VC_270 phage from the *Pseudomonas* P1 strain. This prophage is found in the genomes of many *Pseudomonas* ATUE5 and in some ATUE3 strains. VC_270 seems to be continuously activated, from day 1 post-infection until the last sampling point (Figure 2d). Sporadic activation during the time course could also be detected for VC_400, a widespread prophage in the ATUE5 clade (Figure S7). For all the other prophages, we hypothesize that the right cues for their activation were not present in our controlled environment (such as UV damage or environmental/chemical cues), as it has been previously shown that prophages might be expressed spontaneously at low rates (Nanda et al., 2014, 2015).

The genomes of eleven members of our SynCom encode also tailocins (Table S3, Figure S2), a headless phage tail used for bacterial warfare (Scholl, 2017). All members of the pathogenic community encode a tailocin previously shown to be active against other members of the ATUE5 clade (Backman et al., 2024; Dorosky et al., 2017). The mixed community contained an additional four commensals whose genomes encode for tailocins. When the communities were grown *in vitro*, we saw that the mixed community produced more tailocins than the pathogenic community, as we could not find tailocins in either the pathogenic communities with or without added phage cocktail under the TEM. As expected, bacteriophages were readily seen under the TEM in the community where a phage cocktail had been added (Figure S8).

We carried out plate assays to determine how effective the tailocins encoded by SynCom members were against any of the other SynCom members. We found no evidence for tailocin susceptibility in this specific SynCom. Tailocins are being expressed in the liquid cultures we used to infect the plants, and continue to be expressed during the time course in the plant (Figure 2e), consistent with the stochastic expression of tailocins (Sigal et al., 2024). Tailocins are expressed most highly during the early stages of plant colonization (days 0-2 post infection). From day 5 on, expression remains relatively stable at a basal level, until days 21-26, when the bacterial population drops significantly. That tailocin expression is detectable *in planta* is compatible with an active role of tailocins in shaping bacterial community composition.

### Co-existence of phages and their bacterial hosts in planta

The phages used for the *in planta* experiments can infect most of the bacterial strains in the SyncCom (Figure 3a). We speculated that the introduction of these lytic phages will shift the SynCom composition, putting the most susceptible strains at a disadvantage. Therefore, we used the barcodes to track the abundances of each bacterial strain (Figures 3b, S9, S10). Overall, there were no significant differences in community composition between treatments over the time course of the experiment. Alpha and beta diversity across treatments (Figure 3c, Table S13) did not significantly differ at any time point, either when calculated separately for each replicate or with all replicates pooled. However, specific strains seemed to be affected by the addition of phages: *P. korensis* C3, which is highly susceptible to the PhC91 and PhC95 phages *in vitro* (Figure 1c), had a higher peak abundance in the absence of phages (MC), although this strain seemed to benefit at earlier time points from the presence of phages. A similar trend was apparent for *P. fluorescens* C7, a strain that is weakly susceptible to PhC91, and for *P. viridiflava* P2, which is weakly susceptible to several phages. (Figure 3d and Figure S11). On the other hand, some *Pseudomonas* strains seemed to profit from the reduced growth of the susceptible strains. *P. mandelii* C1, which grows very poorly *in vitro*, reached higher abundances in the presence of phages than in their absence, indicating that it can be competitive *in planta*.

**Figure 3.**
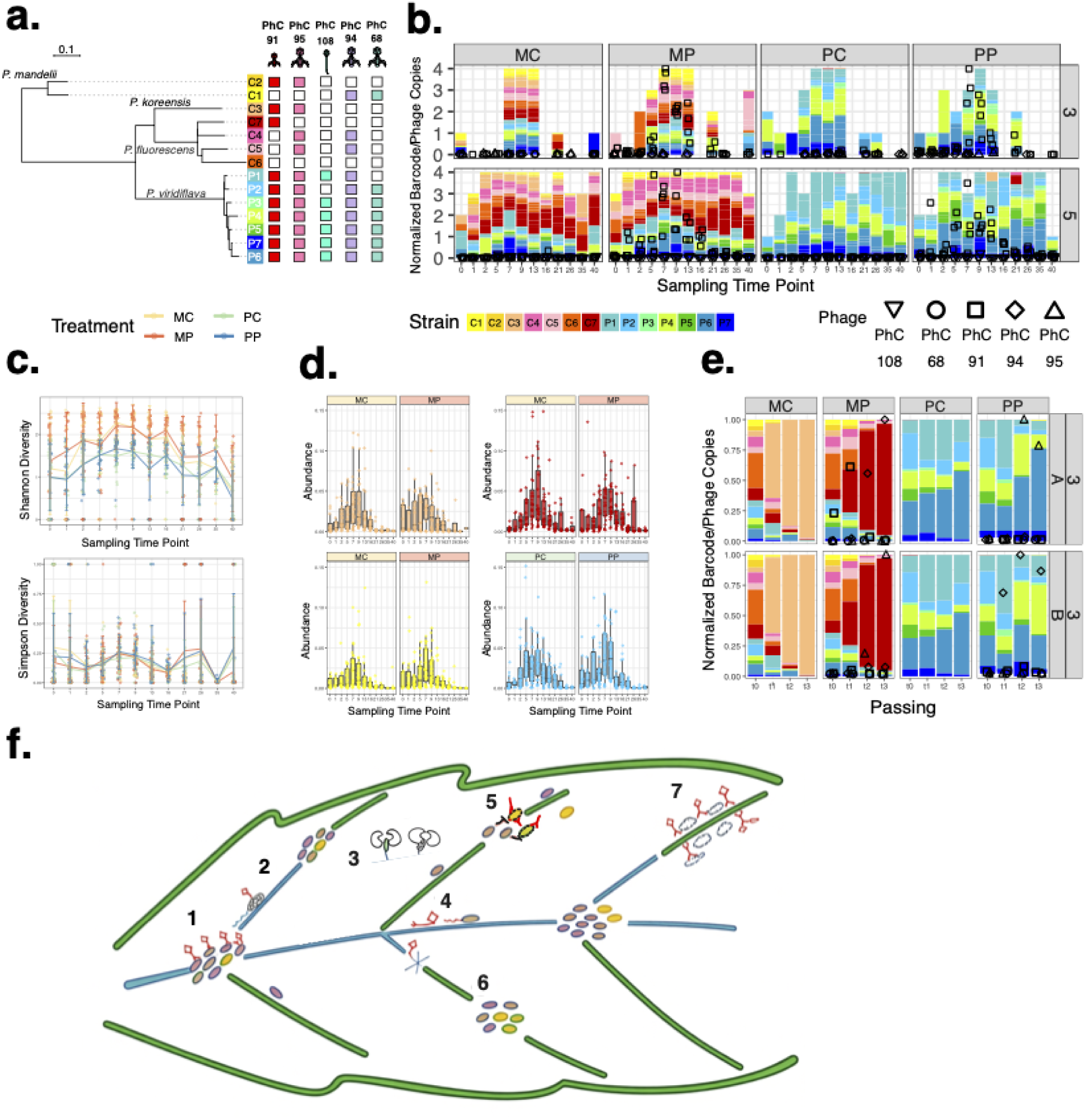
Relative abundance of bacterial strains *in planta* and *in vitro*. **a.** Host range of the phages used in the phage-cocktail against the SynCom members. Colored squares indicate susceptible strains according to the phage color code in Figure 1; white squares denote resistance. Phylogenetic tree of the SynCom members from (Shalev, Karasov, et al., 2022). **b.** Relative abundance of the barcoded *Pseudomonas* strains *in planta*. At each time point, four plants were sampled and used for amplicon sequencing. Missing samples are ones that failed to yield reads after two sequencing runs. Symbols represent bacteriophage abundance calculated by qPCR and normalized to 4 plants per experiment. Only replicates 3 and 5 are shown for clarity; graphs for all replicates can be found in Figure S10 **c.** Alpha diversity as measured by Shannon and Simpson indexes, calculated for pooled samples of each treatment at each time point (n = 1272 samples). Data can be found in Table S13 **d.** Absolute abundance of SynCom members C3, C7, C1 and P2 over the time course of the *in planta* experiments. Absolute abundance was calculated by multiplying the relative abundance of the specific strain by the qPCR value of the barcode. n = 1,272. MC – Mixed community, MP – Mixed community with phages, PC – Pathogenic community, PP – Pathogenic community with phages. For graphs of all strains see Figure S11. **e.** Relative abundance of the barcoded *Pseudomonas* strains in *in vitro* cultures of the SynComs. Symbols denote bacteriophage abundance measured by qPCR and normalized to the maximum per community and experiment. Key to shapes on top. Non-normalized values for phage copies can be found in Table S14, and for the barcode copies of the bacteria in Table S15. **f.** Model for interactions between bacteria and phages on the leaf surface and during subsequent colonization of the leaf apoplast. (1) High humidity enables the creation of a bacteria and phage hot-spot, which facilitates infection. (2) Bacteria infected with milder/slower lytic phages move towards a new niche during the latent phase, effectively spreading the phage. (3) Bacteria can enter the apoplast via open stomata, and infected bacteria can carry the phages to the endophytic communities. (4) Bacteria and phages can use water films to reach novel niches. If the moisture decreases, phages might become trapped and be exposed to abiotic stress. (5) Tailocins are being used by bacteria for bacterial warfare during the colonization stage. (6) Isolated niches are refuges for susceptible bacteria from phages and hostile bacterial strains. (7) Virulent and fast lysing phages rapidly kill all the available hosts in a niche and are now trapped in cell debris. The model was hand-drawn and re-touched with AI.

No strain took over or became extinct during any of the *in planta* experiments, similarly to the microbiomes in the wild (Karasov et al., 2018). When the same SynComs were grown *in vitro*, in a rich, uniform environment, there was a clear hierarchy, with *P. korensis* C3 dominating the MC community in the absence of phages (15/18 cultures), and *P. fluorescens* C7 taking over each replicate in MP communities (Figures 3e, S12). In contrast, in the pathogenic communities PC and PP, all strains co-existed over the passagings, maybe because all phages in the phage-cocktail can reproduce to at least some extent in all PC and PP members, exerting similar pressures on these SynCom members (Figure 3a). However, small and initially undetectable contamination by *P. fluorescens* C5 or C7 led to a rapid and complete takeover by the commensal strains, hinting at a potentially substantial growth advantage that could explain the protective phenotype of this SynCom (Shalev, Ashkenazy, et al., 2022; Shalev, Karasov, et al., 2022) and highlighting the importance of higher-order interactions (Figure S12).

*Pseudomonas koreensis* C3, the highly susceptible strain, behaved in a unique way. When infected with the PhC91 phage *in vitro*, the C3 culture collapsed immediately, never recovering or producing resistant bacteria (Figure 1c, S1). The results were similar in the *in vitro* SynCom context (Figure 3e, S12). *In planta*, a more oligotrophic and fragmented environment, C3 was detectable over the entire length of the plant life cycle; however, when the initial virus/bacteria ratio was high the relative abundance of C3 was lower than that of other susceptible, slower growing bacterial strains, such as *P. viridiflava* P1. This phenotype agrees with pairwise predictive models where high phage density leads to rapid decline of a bacterial population until a steady state is reached (Dey et al., 2026). A possible explanation could be provided by lytic phages that can infect various SynCom members, including fast and slow growers, leading to an “Eliminate the Winner” scenario (Marantos et al., 2022), where the slower growing bacteria have an advantage over the fast growing ones by being less susceptible to the phage, as seemed to be the case for our SynCom when exposed to PhC91 phage *in planta* (Figure 1c). These results lead us to propose a model for phage-bacteria interactions in the leaf (Figure 3f). In this scenario, bacterial hubs are sparsely found along the leaf surface, the presence of a phage in these communities could dictate their fate, but the low connectivity between communities allows other bacterial hubs to grow independently. Water “highways” can connect spatially separated niches, enabling phage particles to infect new hosts and to change the equilibrium at the plant level.

### Bacteria have greater dynamic behavior than their bacteriophages in wild A. thaliana populations

The behavior of our SynCom in the laboratory suggested that the abundance of phages reflects primarily the abundance of their hosts, i.e., that the phages are followers of their hosts. However, a controlled laboratory environment might not reflect the situation in nature. Therefore, we monitored two local *A. thaliana* populations in southwest Germany over their growing season (see Table S1 for the coordinates). We sampled 10-15 plants from each site from October 2024 to April 2025 in monthly intervals, measured bacterial and bacteriophage abundance by qPCR, and analyzed bacterial diversity using 16S rDNA amplicon sequencing. Additionally, several plants from each site were pooled and their DNA was shotgun sequenced to identify bacteriophages as described above. For each plant, we measured the green area, number of leaves, and assessed its developmental stage (Table S16). Local *A. thaliana* germinate in October and reach maturity by the end of February (Figure S13a). Bacterial abundance (measured by 16S rDNA copies) increased with the developmental stage, with a slight drop in mature plants (Figure S13b). As expected, the number of leaves per plant increased early on and then remained stable, (Figure 4a, S13c), while bacterial load increased slightly in plants with more leaves (Figure S13d). Bacterial abundance showed a two-step growth curve over the season, with a first marked increase by mid-autumn, followed by a period of slower reproduction during winter before a further substantial increase in spring (Figure 4b). This is different from what we saw in the laboratory, where bacterial load decreased as plants transitioned to the reproductive phase (Figure 2b). In natural settings, environmental perturbations such as watering events can enable secondary colonizations and facilitate movement within the same or between different leaves, increasing the available niches, and mixing the populations of bacterial commensals, competitors and their phages. Additionally, light hours in our experiment were kept fixed at eight hours a day, which is shorter than what wild plants experience by the end of February in southwest Germany and which might also change microbial dynamics.

**Figure 4.**
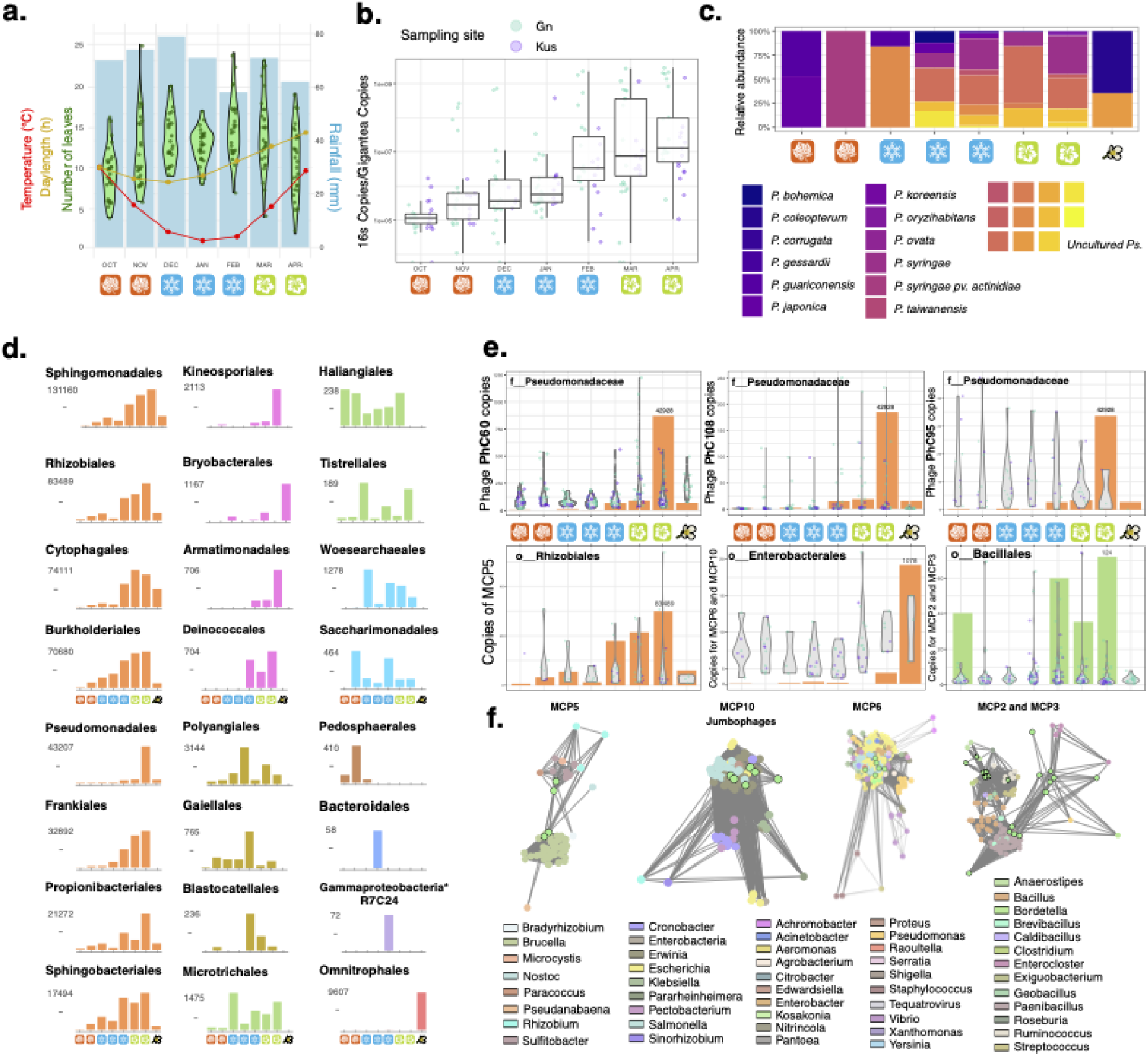
Bacteria and bacteriophages in wild plant microbiomes. **a.** Average temperature (°C, red line), light hours (yellow line) and average rainfall (mm, blue bars) over the *A. thaliana* growing season in sampling sites around Tübingen, Germany. Leaves (green violin plots) were counted for 200 plants sampled in this study. Developmental and morphological attributes can be found in Table S16. **b.** Bacterial abundance over the growing season, calculated by normalizing copies of bacterial 16S rDNA and host *GIGANTEA* gene measured by qPCR. Dots represent individual plants, colored by population. Y-axis is presented in log scale for clarity. **c.** Relative abundance of *Pseudomonas sp.* over the growing season in wild populations of *A. thaliana*, based on 16S rDNA amplicon sequence. **d.** Abundance of different bacterial orders over the growing season. Relative abundance of bacterial strains was calculated by 16S rDNA amplicon sequencing, and then multiplied by 16S rDNA copy number (qPCR) in each plant to obtain the proportion for each bacterial order per plant. Graphs are colored based on similar dynamics during the season. The full graph with all detected orders can be found in Figure S14. **e.** Abundance of isolated and environmental phages and their predicted hosts over the growing season. The abundance of three *Pseudomonas* phages from our collection (PhC60, PhC108 and PhC95) was measured by qPCR of the major capsid protein gene (MCP) and superimposed with the abundance of the Pseudomonads. Metagenomically-assembled phages were clustered according to their gene content, and degenerate primers for the conserved regions of MCP genes of similar phages were used to measure their abundances by qPCR. Hosts were predicted either by the iPHoP tool or by their proximity to known phages in the gene network, as seen in panel f. **f.** One-edge distance gene network of metagenomically assembled (complete and high-quality) phages detected in the wild population samples. Environmental phages are marked in green with a black outline; for the MCP2 and MCP3 network, MCP2 phages are marked with a dotted outline. The other culturable phages are colored according to the legend.

*Pseudomonas* was the fifth most abundant bacterial taxon in wild plants, present during the entire season and peaking by late spring (Figure 4d). While early in the season, the *Pseudomonas* population was dominated by one or two species, it diversified towards spring (Figure 4c). Normalization of 16S rDNA qPCR values by the relative abundance of each bacterial order revealed specific patterns over the season. *Pseudomonas,* similarly to members of other abundant bacterial orders such as Sphingomonadales, Rhizobiales and Burkholderiales, was always present, with increasing abundance as the season advanced (31/120 orders detected, colored orange). Representative seasonal patterns are shown in Figure 4d; see Figure S14 for all bacterial orders. Rarer bacteria showed more diverse seasonal patterns. Of 120 orders identified in our material, only seven (colored pink) also increased in abundance over the season, but with sporadic presence. Thirteen peaked in winter (colored gold), while eleven were least abundant in winter (colored green). Another seven decreased in abundance over the season (colored cyan). The remaining 50 were found sporadically. Some bacterial orders, including Enterobacterales, Staphylococcales, and Omnitrophales, were highly enriched in flowers (Figure S14). Our results contrast with previous findings, where only a handful of bacterial orders showed marked seasonal patterns (Almario et al., 2022). We conclude that sampling time, even within the same season, can greatly influence the diversity and abundance of detected microbes.

To relate bacterial to phage abundance, we focused on phages predicted to be lytic and to which we could assign a possible host, either by host-prediction tools (Table S7) or by gene content similarity (Figure 4f). Based on qPCR results, three phages from our *Pseudomonas*-infecting collection, PhC60, PhC108 and PhC95, could be detected in wild plants (Figure 4e). PhC60 was abundant at all sampling times, even when *Pseudomonas* strains were low in abundance and diversity. PhC108 and PhC95 were present throughout the season, but they were more than an order of magnitude rarer than PhC60. We do not know whether this difference is due to PhC60 being a less effective predator in the laboratory, as it did not completely lyse any culture in our collection (Figure 1b). It has been proposed that being a milder phage could be beneficial when host densities are low and the environment is fragmented (Weitz & Dushoff, 2008).

Other phages similar to Enterobacterales-infecting phages were found at most sampling times (Figure 4e). A correlation between abundance of phages and their hosts becomes apparent when the flower samples are ignored. Phages likely to infect Bacillales were also rare but found in many samples, despite the sporadic presence of their apparent hosts in our material, which may be indicative of a broader host range, or persistence of phage particles in the absence of their hosts.

## Discussion

Bacteriophages must be capable of directly and indirectly affecting microbiomes, yet over the last decade, it has become clear that it is difficult to extrapolate from laboratory assays to the effects of phages on natural microbial communities (Hernandez & Koskella, 2019; Koskella et al., 2022; Papp-Rupar et al., 2025; Svircev et al., 2018). To bridge the laboratory-to-field gap (Lundberg et al., 2025), we compared an ecologically relevant bacterial community and their phages on *A. thaliana* plants in the laboratory and in the wild.

As in many other settings, bacteriophages are abundant in the plant phyllosphere (Dougherty et al., 2023; Forero-Junco et al., 2022; Hulin et al., 2023; Korniienko et al., 2022; Koskella, 2013; Koskella et al., 2011). A first major finding of our work is that the temporal dynamics of bacterial orders were much more pronounced than that of the phages predicted to infect them. This included sampling time points where the predicted hosts could not be detected at all. This is in stark contrast to previous findings: When Rapp-Rupar and colleagues (Papp-Rupar et al., 2025) applied phages to cherry trees in an orchard by spraying and followed their survival over the season, they found that the phages thrived and persisted in the plants as long as the “direct” hosts of the phages were present. This is reminiscent of our findings in controlled laboratory conditions, but differs from our observations in the wild. We propose that some of our findings could be explained both by underestimating a phage’s host range in the wild based on laboratory experiments, and by the ability of phages to persist in the plant environment over seasonal time scales. Specifically, the phages that are the most successful ones *in vitro* or *in planta* in the laboratory might not be the ones most competitive in wild plants. As seen from the gene-content network (Figure S3), most of the phyllosphere-associated bacteriophages are still uncultured and have no isolated relatives, and more metagenomic studies coupled with efforts to isolate and characterize phages are needed to better connect laboratory with field behavior. One limitation of the qPCR method for measuring abundance is that it could amplify DNA of organisms that are no longer alive. To minimize this, we washed leaves sampled from the wild thoroughly before DNA extraction. In addition, we expect the UV exposed leaf microbiome to have a relatively low abundance of intact DNA associated with dead bacteria, so-called relic DNA. Together with the fact that the absolute abundance of phages (and bacteria) did not simply increase over the season, as might be expected for accumulation of relic DNA suggests that we are mostly measuring DNA of members of the plant phyllosphere relevant at the time of sampling.

We find that bacteriophages infecting *Pseudomonas,* including the phages from our collection, are abundant in the phyllosphere. The abundance in the wild of phages characterized in the laboratory greatly varies, with PhC95, the most virulent phage being two orders of magnitude less abundant than the mild phage PhC60. Although it has a broad host range in the laboratory, as a jumbo phage, PhC95 might be more choosy in nature. Such a scenario is consistent with the gene network analysis, which showed its gene *repertoire* to be closely connected only to other local plant-associated phages (Figure 1e). In contrast, PhC60, a small Nickievirus siphophage with a 111 kb genome, is the only phage among our isolates that did not cause complete lysis of any culture in our *Pseudomonas* collection (Figure 1b, S1). PhC91 can lyse many *Pseudomonas* strains in the laboratory, including having a large effect on our *Pseudomonas* SynComs *in planta* and it was isolated from multiple wild plant extracts, but it was not detectable in our plant samples from the wild. These findings agree with general principles of phage biology, where being a fast-reproducing, virulent phage might be counterproductive in the wild. At the same time, in the long-term, being a milder phage enables more flexibility in a changing, fractured environment (Paepe & Taddei, 2006; Roughgarden, 2026; Roychoudhury et al., 2014; Smith et al., 2026).

Spatial scales might be especially relevant in the highly fragmented leaf ecosystem, as spatial structure allows strains to be sheltered from competitors or predators and thus to thrive in certain sub-communities. This might be accentuated in the laboratory, where rain and condensation along with other motile organisms are less of a factor blending different niches (Figure 3f) (Esser et al., 2015; Monier et al., 2009; Saarenpää et al., 2024). Bacteriophages can hitchhike on nematodes in soil (van Sluijs et al., 2025) and rain episodes can lead to an increase in phage diversity along with a decrease of bacterial diversity in soil (Santos-Medellín et al., 2023; Šťovíček et al., 2017). *Pseudomonas* bacteria on their own might not be travelling far on and in a leaf, but rather use “water highways” to physically move and send as well as sense cues for bacteriophage activation (Doan et al., 2020). The diverse landscape of the leaf, including wet and dry, exposed and occluded areas resemble the skin environment, where only some phages co-occur with their hosts, and the composition of bacterial and phage communities depends highly on local abiotic factors (Hannigan et al., 2015).

The makeup of our *Pseudomonas* SynCom greatly changed in the presence of phages in a well-mixed *in vitro* environment, but not when it colonized a leaf. So why does the SynCom become resilient to phage infection in the plant environment? How much is due to altered bacterial physiology in the plant, such as expression of phage receptors, and how much is due to the external and internal topography of the plant leaf? In the wild, we observed an increase in diversity in the *Pseudomonas* community over the season. The higher diversity of potential hosts could be a partial explanation for phages being more easily detectable in the wild than in the phage-treated laboratory samples (Figure 4c,e). While the phages we found can lead a lytic life cycle, prophages are numerically more abundant and could become prominent members of the phage community upon activation, similar to what has been reported for gut microbiomes. Lysogenic phages in the gut are continually being activated at low levels (Sutcliffe et al., 2023), with further enhancement by immune-related cues (reviewed in (Kurilovich & Geva-Zatorsky, 2025)). Many of our metagenome phages cluster tightly with prophages in our *Pseudomonas* SynCom (Figure S3), they encode for genes that enable recombination/integration, and two were integrated into bacterial genomes (Table S8), suggesting that lysogeny is a significant life-style also in the phyllosphere. At this point we can only speculate regarding the cues for their activation and the extent of their influence in these complex bacterial communities.

While our work has raised more questions than it has provided answers, we expect our highly complementary approach that combines laboratory experiments with field observation to inspire future research on plant-bacteria-phage interactions. Questions that need to be addressed include the entire range of phages that a bacterial strain can be infected with, the potential existence of trade-offs between a bacterium’s ability to colonize plants, to compete with other bacteria and to evade phage predation, and how plant genotype and the environment modulate bacteria-phage interactions.

## Materials and Methods

### Pseudomonas cultures and synthetic communities

*Pseudomonas* strains were from a previously described collection (Karasov et al., 2018; Shalev, Karasov, et al., 2022). These were grown overnight at 28°C in LB medium with 10 mg ml^−1^ nitrofurantoin (LB-NF) in a shaker incubator. Overnight cultures were diluted 1:10 and placed back in the shaker for 2.5 – 3 hs, to reach exponential growth phase before use in SynCom experiments. For the SynComs, we diluted the cultures following a published protocol (Shalev, Karasov, et al., 2022). Briefly, the cultures were centrifuged at maximal speed for 10 min, and the pellet was resuspended in 10 mM MgSO_4_. OD_600_ was measured and the density of all cultures was normalized before combining them in equal ratios (Table S11). Strains P1-P7 were used in all four communities (MC, MP, PC, PP), while strains C1-C7 were used for MC and MP communities only. The control treatment was a sterile-filtered 10 mM MgSO_4_ .

### Bacteriophage isolation

*Arabidopsis thaliana* plants from the city center of Tübingen and the outskirts of Gniebl and Kusterdingen (see Table S1 for coordinates) were blended with buffer (0.1 N NaCl, 8 mM MgSO_4_, 50 mM Tris-HCl - pH 7.5) using a kitchen blender and filtered through a series of nets and filters down to 0.45 µm (Millipore) to obtain a plant filtrate, which was stored in glass bottles at 4°C.

Phages were isolated using two methods: dilution-to-extinction and double-agar-layer cultivation.

Dilution-to-extinction: Exponentially grown *Pseudomonas* strains were placed in 96-well dishes (Grainer) and serial dilutions of plant filtrate were added at a ratio of 1:3 to a final volume of 120 µl. Dishes were incubated overnight at 28°C and then screened for clearance of wells compared to an untreated control. The content of promising wells was passed through a 0.45 µm filter (Millipore) and used in the double-agar layer method for further purification.

Double-agar-layer: Exponentially grown *Pseudomonas* bacteria were mixed with warm LB-NF-agar containing 5 mM CaCl_2_, 5 mM MgCl_2_ and 0.1 M MnCl_2_ to a final concentration of 0.3% agar. The mixture was plated on top of a regular agar plate and left to dry. For bacteriophage isolation, we followed a published high-throughput protocol (Kauffman & Polz, 2018) with three rounds of purification.

Many bacteriophages were lost during this process, probably due to the rise of resistance in the bacteria. Some bacteriophages described in this study could not be kept viable over time (Table S1).

### TEM imaging

Phage samples were placed onto glow-discharged formvar- and carbon-coated copper grids and negatively stained with 1% uranyl acetate. Images were acquired with a CMOS camera (TemCam-XF416, TVIPS, Gilching, Germany) mounted either on a Tecnai Spirit transmission electron microscope (Thermo Fisher Scientific, Eindhoven, The Netherlands) operated at 120 kV or a JEM-2100Plus (Jeol GmbH, Freising, Germany) operated at 200 kV.

### Nucleic acid extraction

Nucleic acids were extracted from plants following a published protocol (Haeussler et al., 2025) with the following modifications: samples were flash-frozen in tubes containing a mixture of silica and glass beads ranging from 0.1 to 5 mm and stored at -70°C until extraction. Plants were shattered by bead-beating in a Precellys tissue homogenizer (Bertin Technologies) (1 min, 30 sec pause, 1 min). Warm extraction buffer was added, and samples were flash-frozen and placed in -70°C until further treatment. Plants sampled in the wild were bead-beaten twice before adding the extraction buffer and once afterwards. Samples were re-thawed on ice and centrifuged in a table top centrifuge (Eppendorf) for 30 min at 4°C to pellet cell debris. The supernatant was transferred to Econospin (Epoch Life Science) columns for nucleic acid extraction following the protocol in (Haeussler et al., 2025). After elution, samples were divided in two halves each, receiving either 1 µl RNAse-I or 1 unit of Turbo-DNAse (Thermo Scientific), and incubated at 37°C for 30-60 minutes. RNA samples were treated with EDTA (final concentration 15 mM) and heated to 75°C for 10 minutes for DNAse inactivation. All samples were frozen until further use. PCR with primers targeting the *A. thaliana GIGANTEA* gene (Lundberg et al., 2021) was performed on all RNA samples to ensure DNA absence.

DNA of bacteriophages and *In vitro* SynComs was extracted using Wizard columns (Promega) (Poulos & Sullivan, 2016). DNA/RNA concentration of the samples were calculated using the Quant-it PicoGreen and RiboGreen kits (Invitrogen), or measured by Nanodrop (Thermo Scientific). Samples were normalized according to the maximum possible concentration for each experiment (1 µg for RNA samples). For RNA samples, cDNA was synthesized with the LunaScript RT Supermix kit (New England Biolabs). cDNA was diluted to 300 µl with DEPC-water and used as template for qPCR.

### Illumina shotgun sequencing

Libraries for DNA sequencing (isolated bacteriophages, wild plants) were prepared according to an in-house protocol, with DNA being fragmented using bead-bound Tn5 transposomes (Hydrophilic Streptavidin Magnetic Beads, New England Biolabs), and ligated to Nextera (Illumina) adapters for sequencing. Samples were pooled, concentrated and cleaned with SPRI beads at a 1:1 ratio, washed twice with 70% ethanol and eluted in TE. Size selection was done using BluePippin (Sage Science) to select fragments within a 300-700 bp range. DNA concentration was measured using Qubit (dsDNA high sensitivity kit, Thermo Fisher), and the average length of inserts was assessed with a Bioanalyzer2100 (Agilent). Libraries were sequenced using Illumina MiSeq or HiSeq 2000 platforms with 2 x 150 bp reads.

### Genome assembly and annotation

Reads were trimmed using Trim Galore (v0.6.10) (Krueger, 2015) and de-novo assembled with SPAdes (v3.15.5) (Prjibelski et al., 2020) with default parameters for single strains and with MEGAHIT (v1.2.9) (Li et al., 2016) with default environmental parameters for metagenomes. Contigs from metagenomes were further extended using COBRA (v.1.2.3) (Chen & Banfield, 2024). Viral contigs in the assemblies were further identified with geNomad (v1.5.2) (Camargo et al., 2024), and their quality was assessed with CheckV (v1.0.3) (Nayfach, Camargo, et al., 2021). Gene annotation was done with the Bakta pipeline (v1.11.0) (Schwengers et al., 2021). For genes without a specific annotation, a second annotation layer was performed using InterProScan (v5.74.105) against the following databases: Pfam, Panther, Superfamily, Prints, FunFam, CDD (Blum et al., 2025; Jones et al., 2014). Genomes were manually inspected in Geneious Prime (v2025-v2026). Bacterial contigs were further scaffolded using RagTag v 1.0.1 employing the closest closed *Pseudomonas* genome available at NCBI.

### Host range assessment of bacteriophages

To assess the host range of *Pseudomonas*-infecting phages, we used the double-agar layer method for creating plaques (as described above), together with monitoring OD_600_ in liquid cultures. For the latter, we placed 190 µl exponentially growing cultures at an early phase in a 96-well plate. We added 10 µl LB to the uninfected controls and 10 µl phage lysate to the samples. Every experiment was performed in triplicate. Plates were incubated in a TECAN plate reader (Robot Tecan Infinite M200, Tecan Life Sciences) at 28°C over 6-20 hr with the following program: shake (5 sec), rest (5 sec), OD_600_ measurement, idle (4 min).

To determine the host-range of each phage, we classified each growth curve on a scale from 0 to 2, where 0 is no effect of the phage, 1 is attenuated growth after an initial phase that was the same in control and experiment, and 2 is complete lysis. Some susceptible strains were also treated with 10 µl of boiled phage lysate, to assess any possible growth inhibitors in the lysate. No such effects were found.

### In planta infections

*Arabidopsis thaliana* seeds were sterilized by overnight placement at -70°C, followed by at least 4 h of vapor-phase sterilization with chlorine gas. Seeds were stratified at 4°C for 4-6 days and sowed in 195 ml pots on soil (CLT Topferde, einheitserde.de).

Plants were grown at 23°C with 8 hr 125–175 μmol m^-2^ s^-1^ and relative humidity of 65%. Accessions used are detailed in Table S11. On day 20/21 after sowing, plants were thoroughly watered and covered for 2 hr. Plants were separated into five groups in a semi-random manner: groups of five plants similar in size and leaves number were split, one for each treatment, so that at the end of the process all treatments had similar plant profiles.

Plants were each sprayed for 2 sec with the corresponding microbial community (MC, MP, PC, PP, Control) with an airbrush (BADGER 250-1, Badger Air-Brush Co.), and left to dry covered with transparent hard plastic covers for 1-2 hr. Each plant was assigned a “sampling day” and placed in a new tray along all plants from all treatments belonging to the same “sampling day”. After 3 hr, plants for time point 0 (t0) were harvested, and the rest placed back in the growing chamber.

Harvesting was done by cutting the aerial part of the plant, weighing, then placing it in a tube with glass and silica beads followed by flash-freezing. Plants were always sampled at mid-day. Samples were stored at -70°C until the end of the experiment, when all samples were processed together for nucleic acid extraction.

Plants were sampled every two to three days (Table S11), the last day being the time point when the plants had almost open flower buds. Plants too large to be placed in a 2 ml tube were placed in 5 ml Eppendorf tubes or 50 ml Falcon tubes, flash-frozen, crushed with a pistil, and the powder was transferred to a 2 ml Eppendorf tube containing glass and silica beads. Entire plants were collected for each time point (without roots), with four plants per treatment. For visualization purposes, the days of sampling were uniformly categorized for all replicates, so that some sampling time points are shifted by 2 d on the graphs (Table S11).

### In vitro SynCom experiments

For serial transfers, SynComs were diluted into LB-NF medium at a ratio of 1:10. 1 ml was used for DNA extraction using Wizard columns as described above (t0). The rest was incubated in a shaking incubator at 28°C. Cultures were diluted every day in a 1:10 ratio, and DNA was extracted at every time point using the Wizard Columns kit (Promega). A control culture of pure LB-NF was used to monitor for contaminations. DNA was quantified using Nanodrop (Thermo Scientific) and normalized prior to quantification of phages by qPCR and amplicon library preparation for the bacterial barcodes. TEM pictures of the *in vitro* communities were taken for experiment 5, before the daily dilution of time points t1-t3.

### Barcode quantification

Barcodes were quantified by qPCR using the primer pair 79.80 (Table S17) as described below (“qPCR quantification”) for all *in planta* experiments. Despite the control treatment having no added barcoded bacteria, all samples showed a certain level of noise, with the control treatment background being almost constant throughout the course of each experiment. In experiments 1, 3 and 4, where the background has high and the coefficient of variation within the control values was close to 0 (Table S18, Figure S15a,b), we subtracted the average value of the control treatment from the copy number values in experimental samples, so that we could accurately assess barcoded bacterial abundance. Corrected copy numbers that were negative were reset to 0 copies for visualization purposes. Absolute barcode copy numbers were used to calculate the absolute abundance of each strain in each sample by multiplying it by the relative abundance of that strain, as calculated from barcode amplicon sequencing (Table S19).

### Wild plant sampling

*Arabidopsis thaliana* plants were sampled in the outskirts of Gniebl and Kusterdingen, in Baden-Württemberg, Germany (See Table S1 for exact location and coordinates). Plants were sampled at mid-day once at the end of each month between October 2024 and April 2025. Plants were photographed, cut and placed in a tube on ice for up to three hours before further processing. Plants were thoroughly washed with double-distilled water, photographed and flash-frozen in a tube with the same bead mixture as described above. DNA was extracted and normalized as described above. Each photo was inspected to count leaf numbers, determine its developmental stage (juvenile, adult, bolting, flowering,) and to measure with ImageJ (Abramoff et al., 2004) its diameter (Table S16). Environmental data for Figure 4a were obtained from https://en.climate-data.org/europe/germany/baden-wuerttemberg-363/r/october-10/.

### qPCR

DNA and cDNA samples were normalized to the maximum possible concentration per experiment (*in vitro* communities 1-6 / *in planta* infection experiments 1-6 / wild plants). Each DNA sample was split in two, so that each qPCR plate included two replicates per sample. Additionally, each qPCR plate included a set of linearized standards of a specific PCR product in pGEM (Promega). The standard curve was calculated by comparing the Ct cycle for 10^3^ to 10^8^ copies of the amplicon. All reactions for the same experiment used the same sample DNA concentration to enable quantitative comparisons between reactions. qPCR used a Bio-Rad CFX384 Real Time System, with the following cycles: 3 min denaturation at 95°C, 40 cycles of 20 sec denaturation at 95°C, 20 sec annealing at the T_m_ temperature specified in Table S17, and 30 sec elongation at 60°C, ending with a melting curve. At the end of each cycle, fluorescence was measured. 10 µl reactions contained 5 µl of the 2x Luna Universal qPCR master mix (New England Biolabs), 0.25 µM primers, and 2 µl template. Copy numbers for the amplified fragments were calculated based on the standard curve, which had an R^2^ > 0.97. For cDNA samples, the same procedure was used, except with triplicates.

### Amplicon sequencing - Pseudomonas barcodes and 16S rDNA of wild plants

Bacterial barcode sequences were amplified using primers 77.78 with the following parameters: initial denaturation for 3 min at 95°C, followed by 25 cycles of 30 sec denaturation at 95°C, 30 sec annealing at 55°C, and 30 sec elongation at 72°C, with a final 3 min elongation step. Samples were cleaned by adding SPRI beads in a ratio of 0.8:1, incubated for 5 min, and the supernatant was discarded. Beads were washed twice with 200 µl 70% ethanol and left to dry for up to 10 min. DNA was eluted from the beads using the mix of the second PCR, where primers were replaced by Nextera barcode-adapters for Illumina sequencing. Both reactions were performed using Q5 high-fidelity DNA polymerase (New England Biolabs) in 25 µl: 1x Q5 buffer, 0.02 units Q5 polymerase, 0.2 µM primers/adapters, 0.2 µM dNTPs. The second PCR was denatured for 30 sec at 98°C, followed by 8 cycles of 15 sec denaturation at 95°C, 20 sec annealing at 55°C, and 30 sec elongation at 72°C. Samples were pooled at equal volume, cleaned and concentrated using SPRI beads as described above, and eluted in TE water.

Samples with bacterial barcode amplicons were sequenced twice in independent runs with 2x150 bp reads, while the 16S rDNA amplicons were sequenced using 2x300 bp reads on an Illumina HiSeq 2000 platform. Barcode read counts of each run were combined to obtain the total number of reads per strain per sample. Strains with fewer than 20 combined reads were disregarded in further analyses. Samples where the total number of reads from all strains in both sequencing runs amounted to fewer than 55 were also removed. Time points where only one out of four plants for the same treatment and experiment yielded reads were disregarded for the plots. Statistical analyses were performed in R with dplyr (Wickham et al., 2020), tidyr (Wickham & Lin, 2020), stringr (Wickham, 2019), vegan (Oksanen et al., 2020), broom (Robinson & Hayes, 2020), and purr (Wickham & Henry, 2020).

For 16S rDNA amplicons, wild plants were processed as described for laboratory samples, but the first PCR was carried using the 87.88 primer pair. 16S rDNA amplicon annotation was performed using Qiime 2 (Bolyen et al., 2019) against specific databases (Bokulich et al., 2018; Kaehler et al., 2019; Quast et al., 2013; Robeson et al., 2021). Unique 16S rDNA amplicons of *Pseudomonas* were assigned a taxonomy using BLAST (Altschul et al., 1997) against the nr database of NCBI.

### Gene similarity networks

To assess gene content similarity between bacteriophages genomes we used PanGene-O-Meter (Ashkenazy & Weigel, 2025) based on DIAMOND DeepClust (Buchfink et al., 2026) to identify protein clusters sharing sequence identity of at least of 80% and mutual coverage of 70%. Genomes were clustered based on GCSj of 50%. The GCSj distance calculated for each genome-genome comparison was deducted from 1 to obtain a distance metric (instead of similarity) and used in Cytoscape (Shannon et al., 2003) to visualize and color the clusters. Bacteriophage genomes for the network were downloaded from the INPHARED database (Cook et al., 2021). Only dsDNA phages were used for the network of our dsDNA phages, and only ssDNA phages for our Microviridae isolate. For visualization purposes, the networks show only the nodes directly related to our relevant phages (“single-edge distance”), as the number of phage genomes in the databases is too large to be presented in full.

### Host prediction and taxonomy of bacteriophages

To assess the taxonomy of phage isolates and environmental contigs, we used taxMyPhage (Millard et al., 2025) with the VMR_MSL40.v2.20251013 database and default parameters. For the majority of phages/contigs, the only classification obtained was the one from geNomad (Camargo et al., 2024). Host prediction was performed with iPHoP v1.3.2 (Roux et al., 2023) for all “complete” and “high quality” environmental contigs.

## Author Contributions

S.R. and D.W. conceived the project. S.R. designed the experiments. S.R., V.H-W. and C.C. performed experimental work. E.M.H performed image analyses. S.R, V.H-W. and N.B. sampled in the field. K.H. performed TEM imaging. S.R. and H.A. performed data analysis. S.R. and D.W. supervised the project. S.R. drafted the paper, which was critically revised and approved by all authors.

## Data and code availability

Uncropped TEM pictures can be found in the raw data repository. Raw reads for the metagenomic samples, genomes of isolates and metagenomically-assembled phages (complete genomes) have been deposited at ENA, Project PRJEB110834.

## Acknowledgements

We thank R. Schwab, M. Lucke, H. Pfingst and B. Bhawna for help with field work, S. Darwish for help with laboratory work, I. Koch for help with TEM imaging, D. Ibarra and S. Geiger-Rudolph for help with library preparation and sequencing, and I. Brezukov for help with data handling. This work was funded by EMBO post-doctoral fellowship ALTF 445-2023 (S.R.), an Athene grant from the DFG-funded Excellence Cluster “Control of Microorganisms to Fight Infections” (S.R.), the Christiane Nüsslein Volhard Foundation (S.R.), an Erasmus+ scholarship (C.C.), the European Research Council through ERC-SyG PATHOCOM 951444 (D.W.), the DFG-funded Excellence Cluster CMFI (D.W.), the Novozymes Prize of the Novo Nordisk Foundation (D.W.), and the Max Planck Society (D.W.).

