## Supplementary material for "Dynamic co-existence of bacteriophages and their hosts in the *Arabidopsis thaliana* phyllosphere": Figure S

### Supplementary Figures

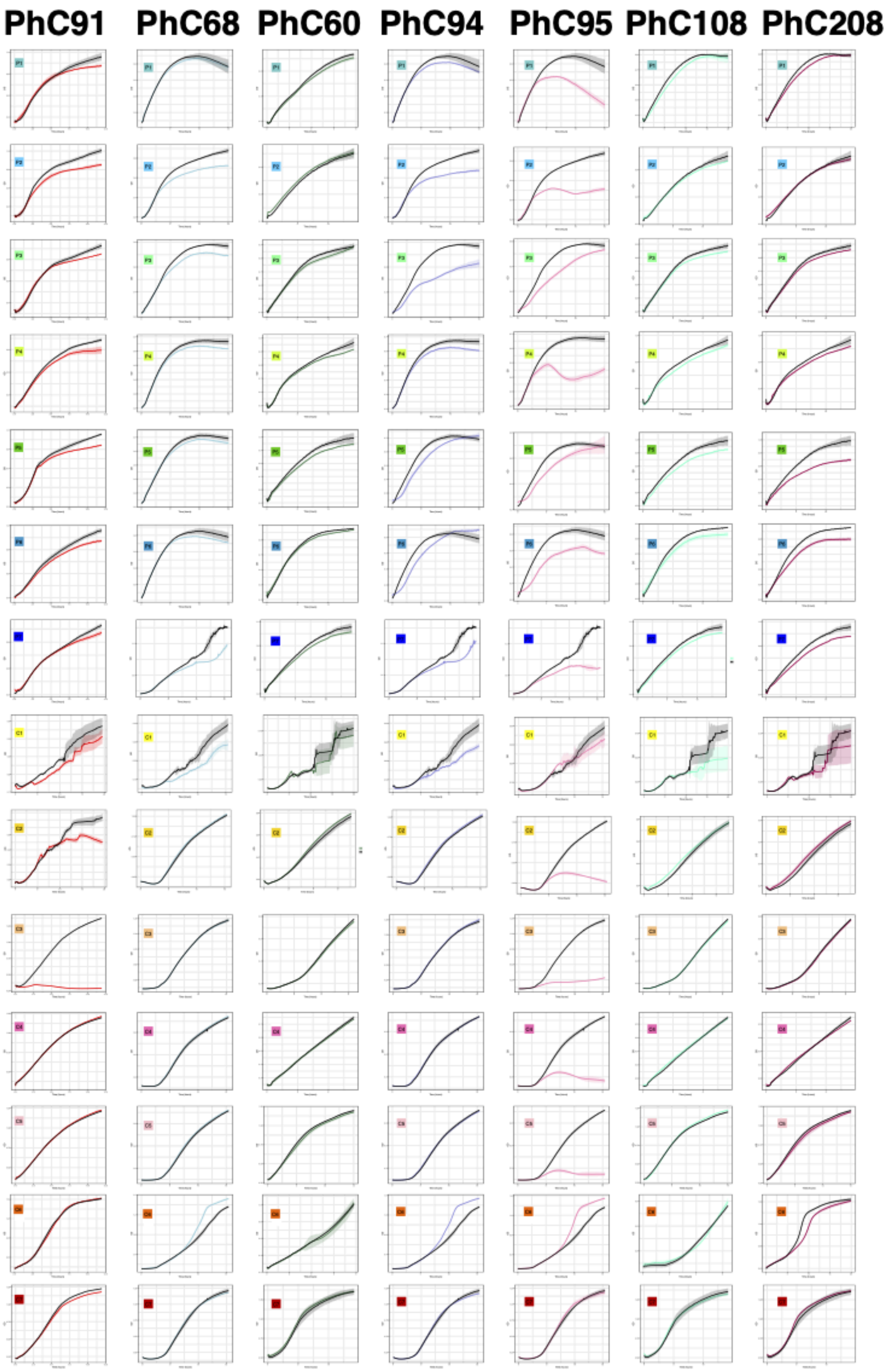


**Figure S1. Host range of isolated phages.** Infection curves as measured in liquid assays over ~15-20 hours of the phages isolated in this study and the members of the *Pseudomonas* SynCom. Cultures were grown and measured in triplicates, SD is denoted with bars. Black dots mark control, uninfected cultures, colored dots are infected according to the legend in Figure 1a at 0.05% v/v. SynCom strains P1-P7 and C1-C7 are noted inside the graphs.

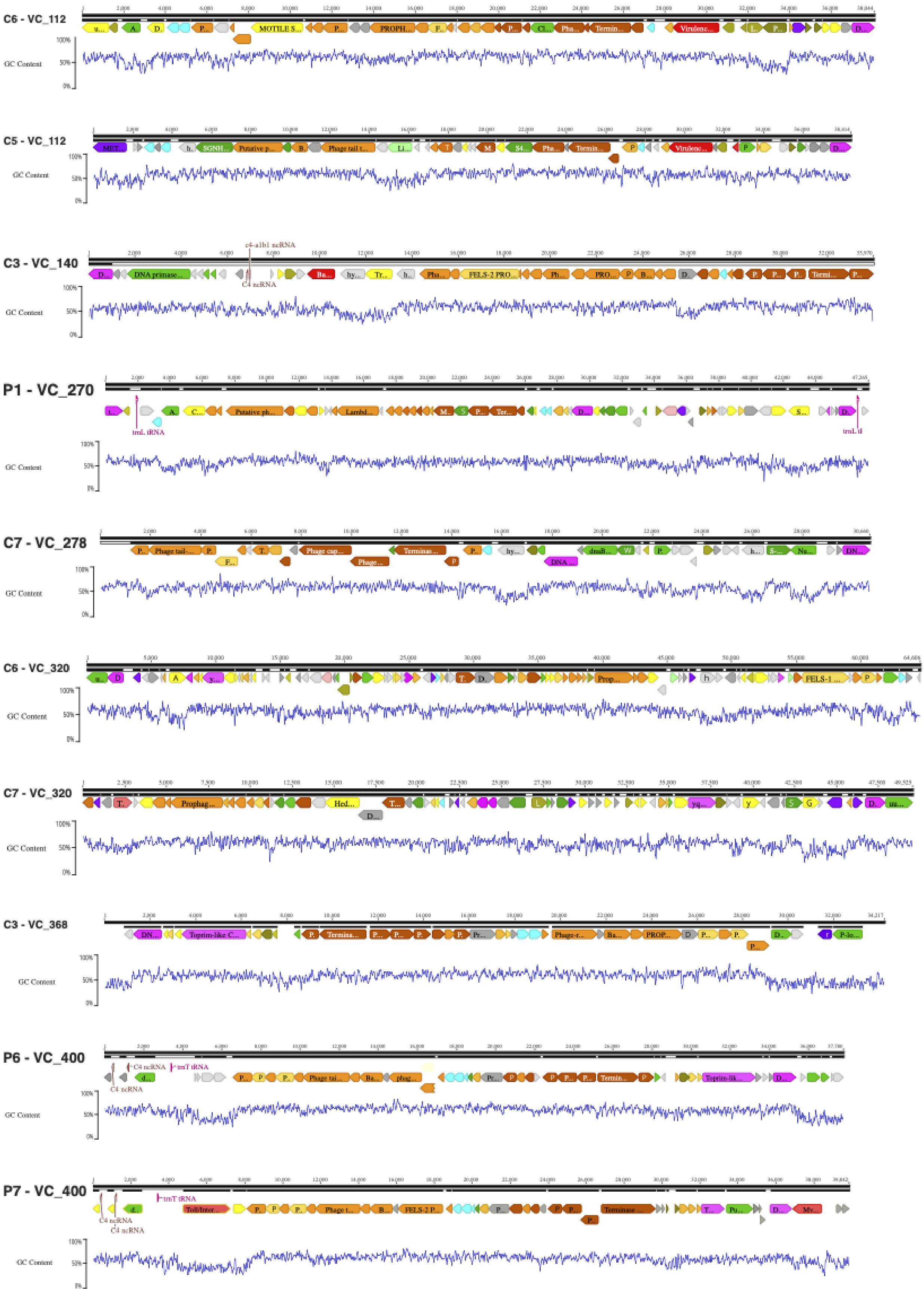
C6 - VC_112
GC Content
C5 - VC_112
GC Content
C3 - VC_140
GC Content
P1 - VC_270
GC Content
C7 - VC_278
GC Content
C6 - VC_320
GC Content
C7 - VC_320
GC Content
C3 - VC_368
GC Content
P6 - VC_400
GC Content
P7 - VC_400
GC Content

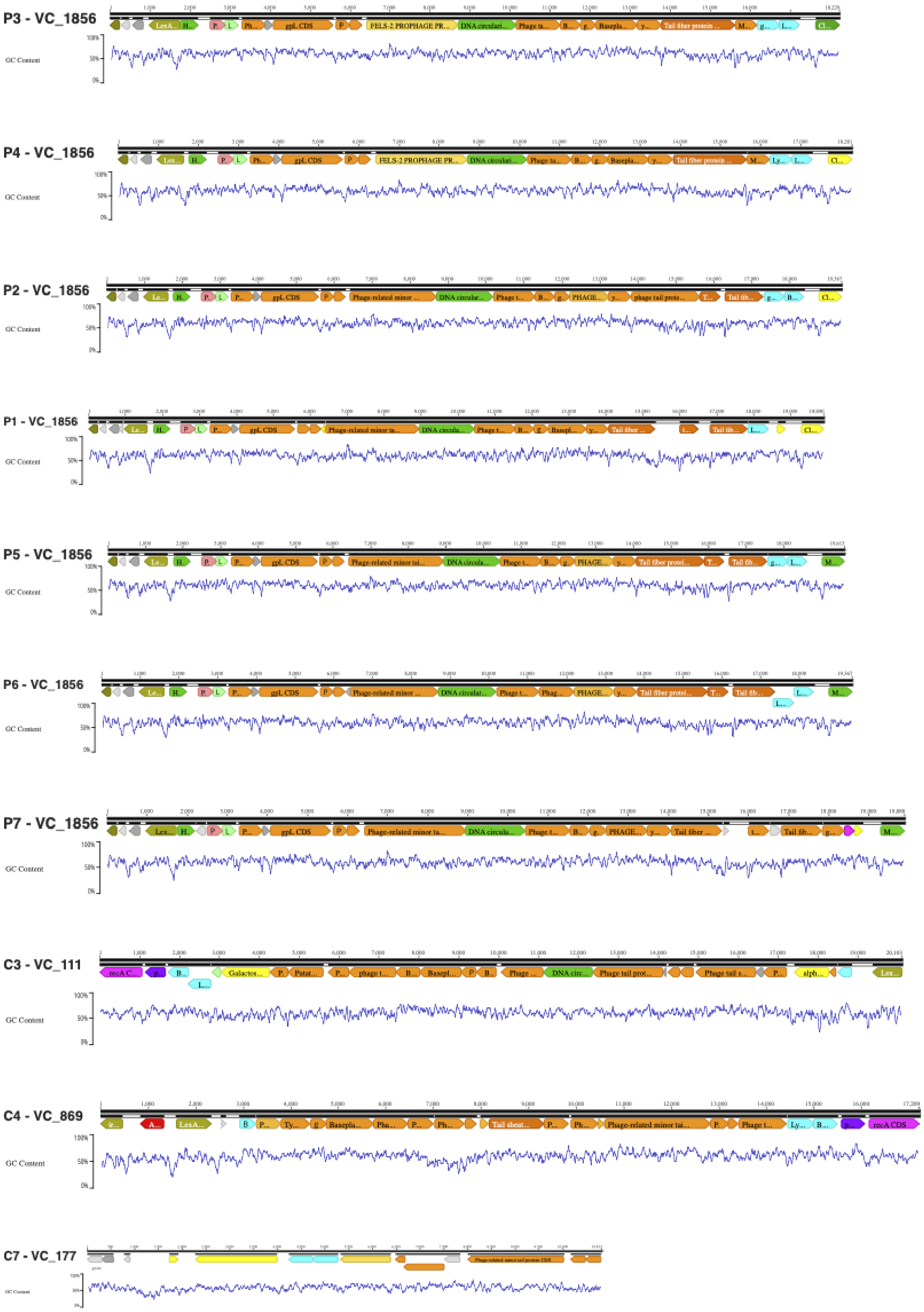
P3 - VC_1856
GC Content
P4 - VC_1856
GC Content
P2 - VC_1856
GC Content
P1 - VC_1856
GC Content
P5 - VC_1856
GC Content
P6 - VC_1856
GC Content
P7 - VC_1856
GC Content
C3 - VC_111
GC Content
C4 - VC_869
GC Content
C7 - VC_177

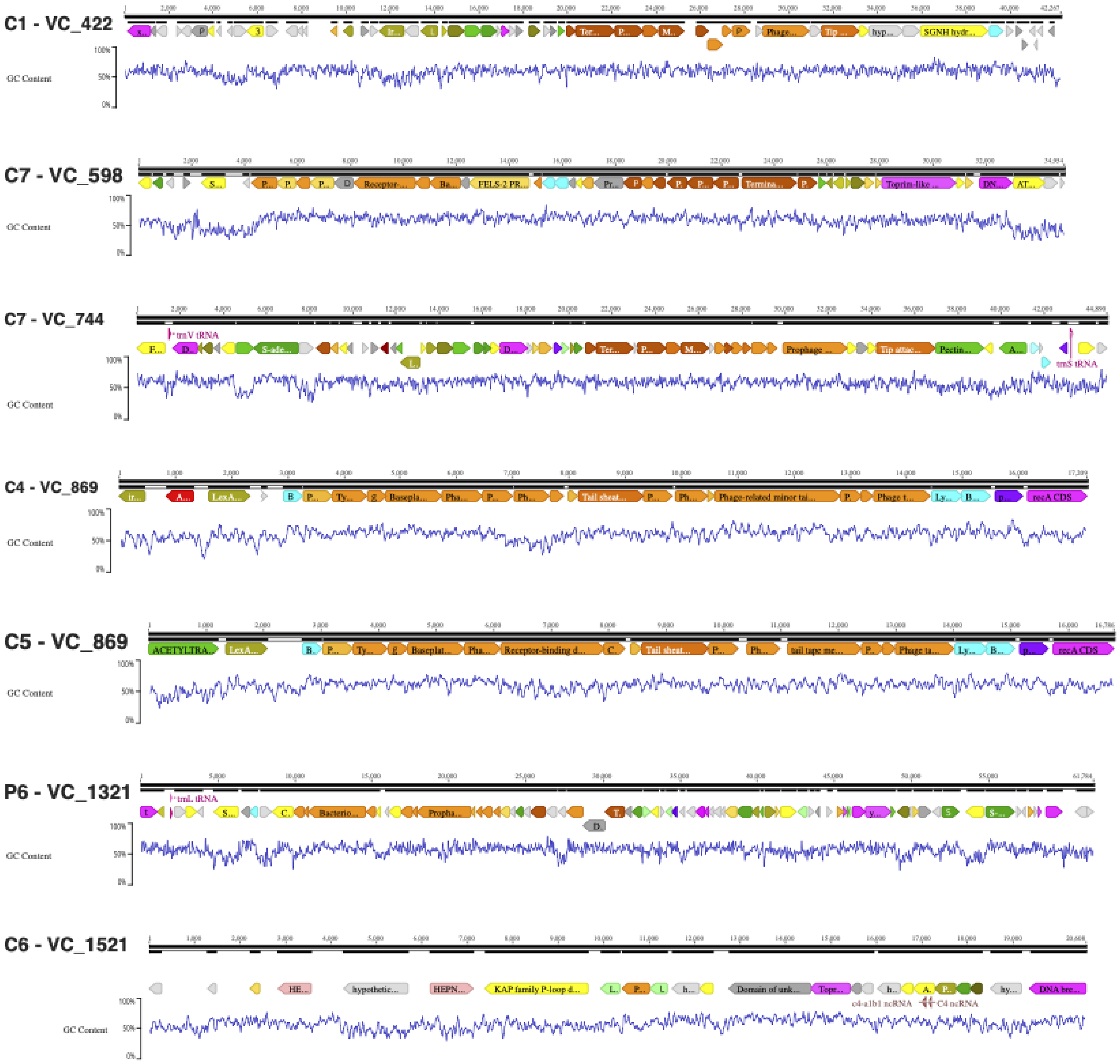


**Figure S2. Genomes of prophages and tailocins encoded by the *Pseudomonas* SynCom.** Schematic representation of the prophages and tailocins, named as the strain of origin and their cluster. Numbers on top denote bp; blue line the GC content. tRNAs are marked in magenta, non-coding RNAs in red. ORFs are colored according to the predicted function: Orange – Structural genes (light - tail-related genes, dark – head related genes); bright green – metabolism and DNA synthesis; red – toxin/antitoxin, defense systems, immunity genes; purple – auxiliary metabolic genes; olive green – regulatory genes; light green – lipoproteins; cyan – lysis-related proteins; dark grey – conserved hypothetical proteins; light grey – hypothetical proteins; yellow – classification unclear.

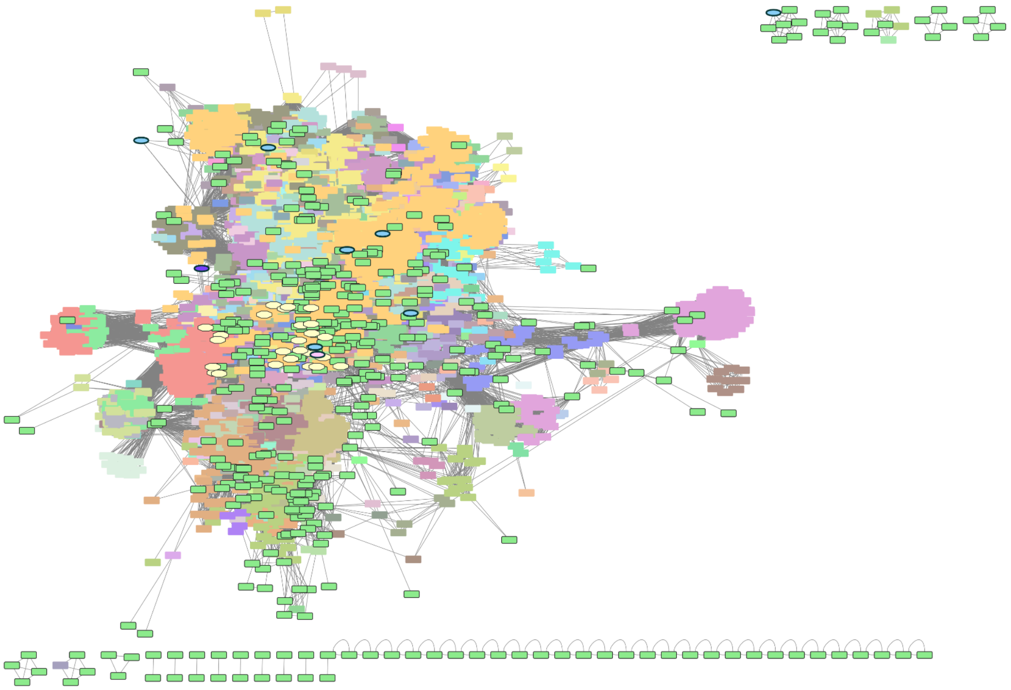

**Figure S3. Gene-to-gene network of dsDNA phages.** dsDNA phages isolated in this study (ellipses colored according to Figure 1a), prophages encoded by the Pseudomonas SynCom (yellow ellipses), and phage genomes assembled by metagenomics in this study (bright green rectangles) were clustered by gene content and similarity with known phages. Only clusters directly connected to the phages relevant to this project are shown. Phages are colored according to their isolation host species, as reported in the INPHARED database (Cook et al., 2021).

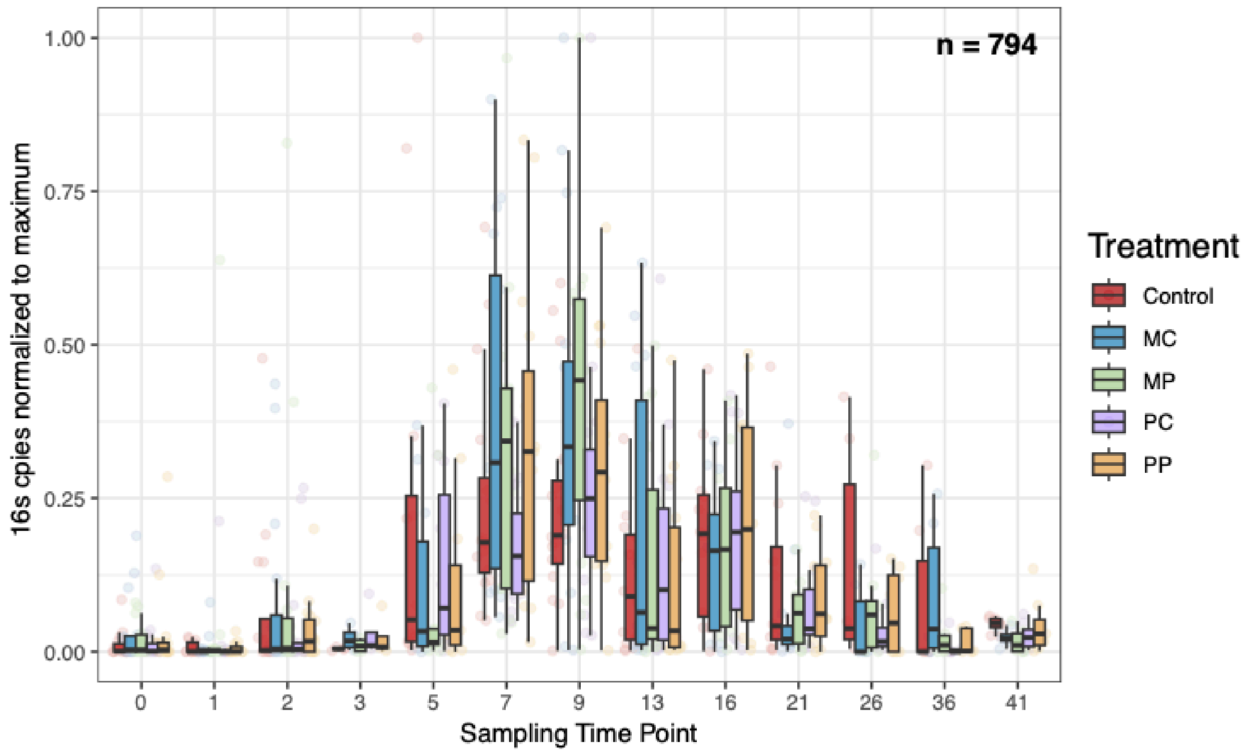


**Figure S4. Bacterial abundance *in planta*.** Abundance of bacteria *in planta* was measured by qPCR on the 16S rDNA gene. For each experiment the copy numbers were normalized to one to facilitate visualization. MC-Mixed community, MP- Mixed community with phages, PC- Pathogenic community, PP- Pathogenic community with phages.

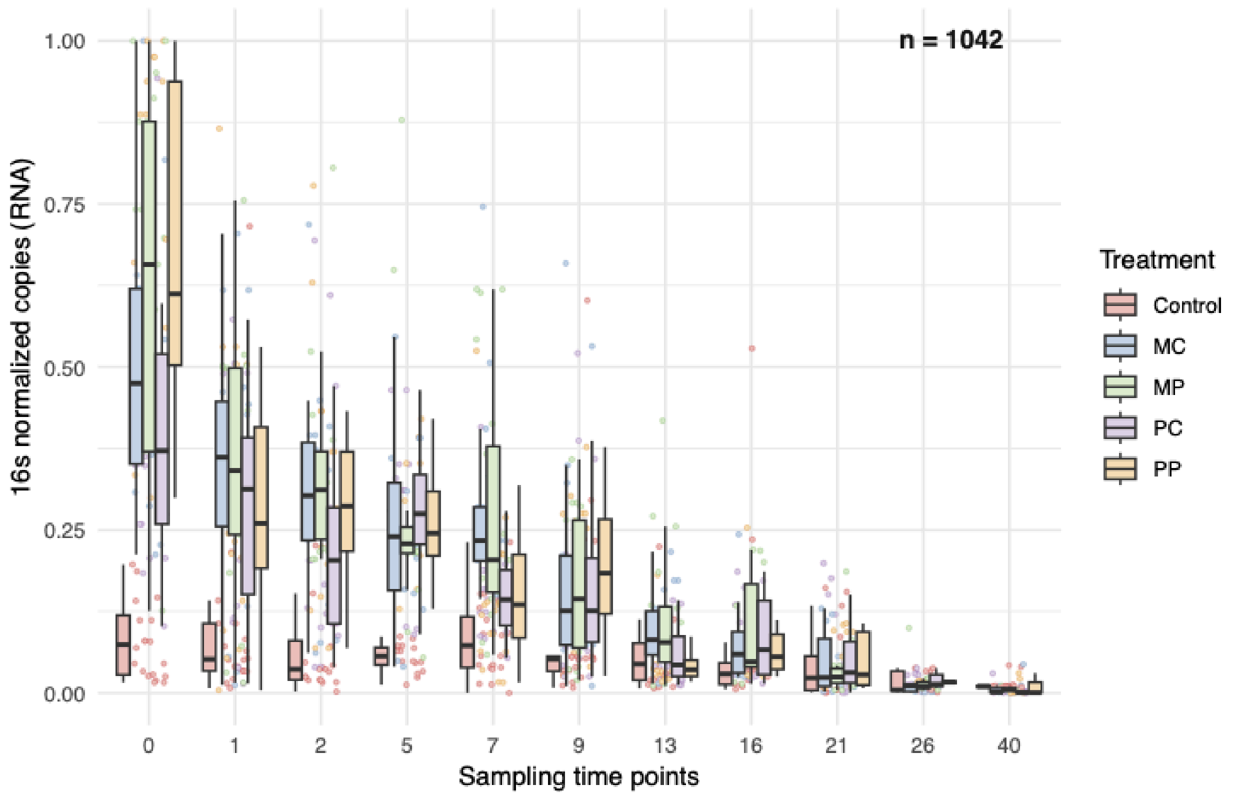


**Figure S5. Bacterial activity *in-planta*.** Bacterial activity was measured by qPCR on the 16S rRNA transcript. Copies were normalized to one per experiment to enable visualization. MC-Mixed community, MP- Mixed community with phages, PC- Pathogenic community, PP- Pathogenic community with phages.

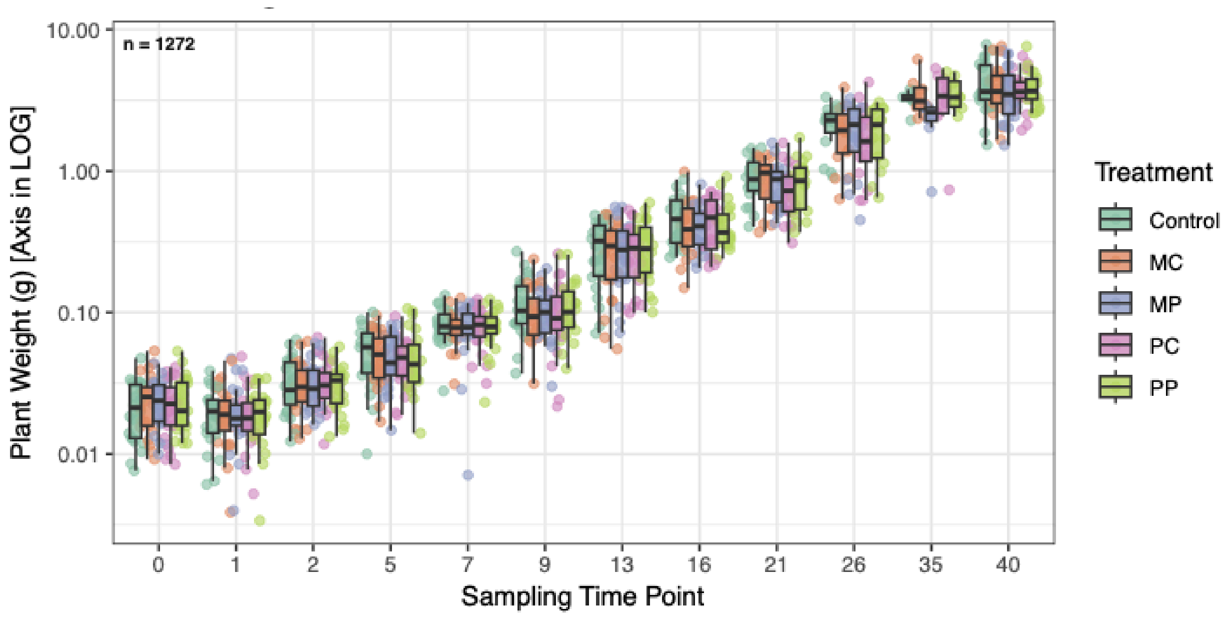


**Figure S6. Plant weights.** Plant weights (grams) at the time of sampling, per treatment. Sampling time points are post-infection, so that 0 represents 21-22 days old plants. Y-axis is presented in log scale to facilitate visualization. 1272 plants were measured in 6 experimental replicates.

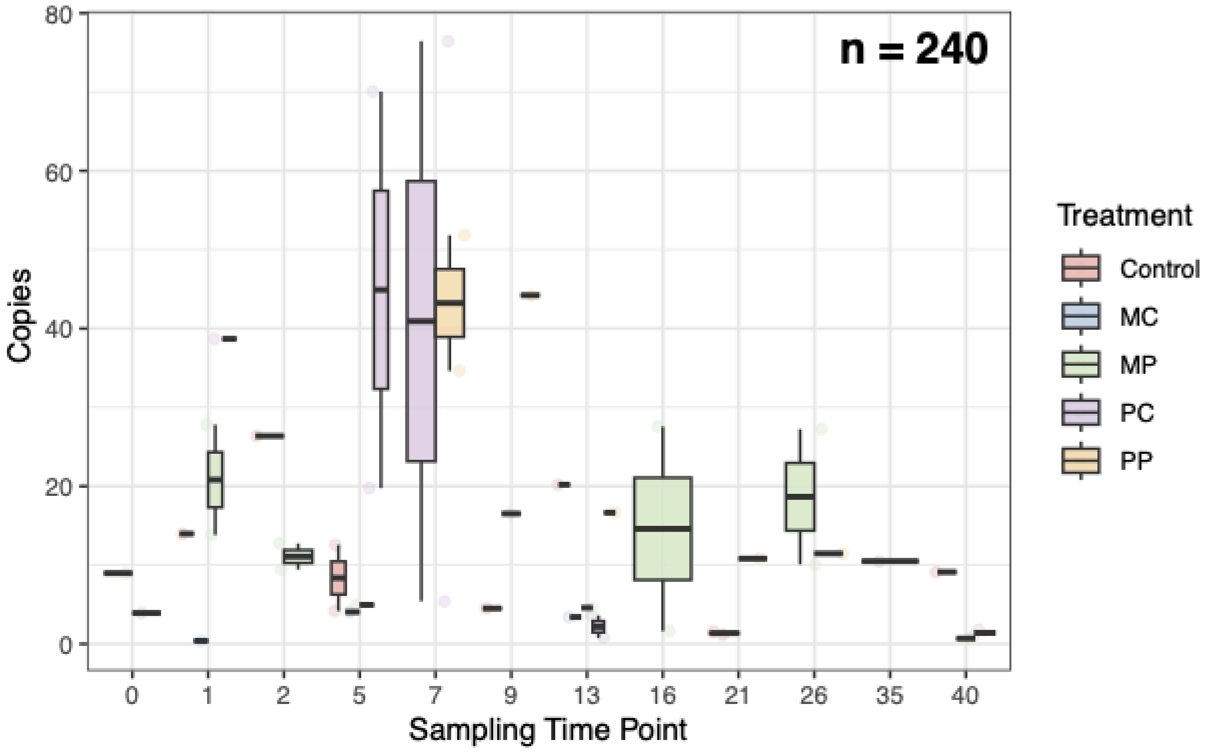


**Figure S7. Prophage VC_400 expression *in planta*.** Expression pattern of a *P. viridiflava* prophage. The expression was measured by qPCR on a structural gene of the prophage (primers 63.64) and the copy number was calculated by deducting the background noise. n = 240

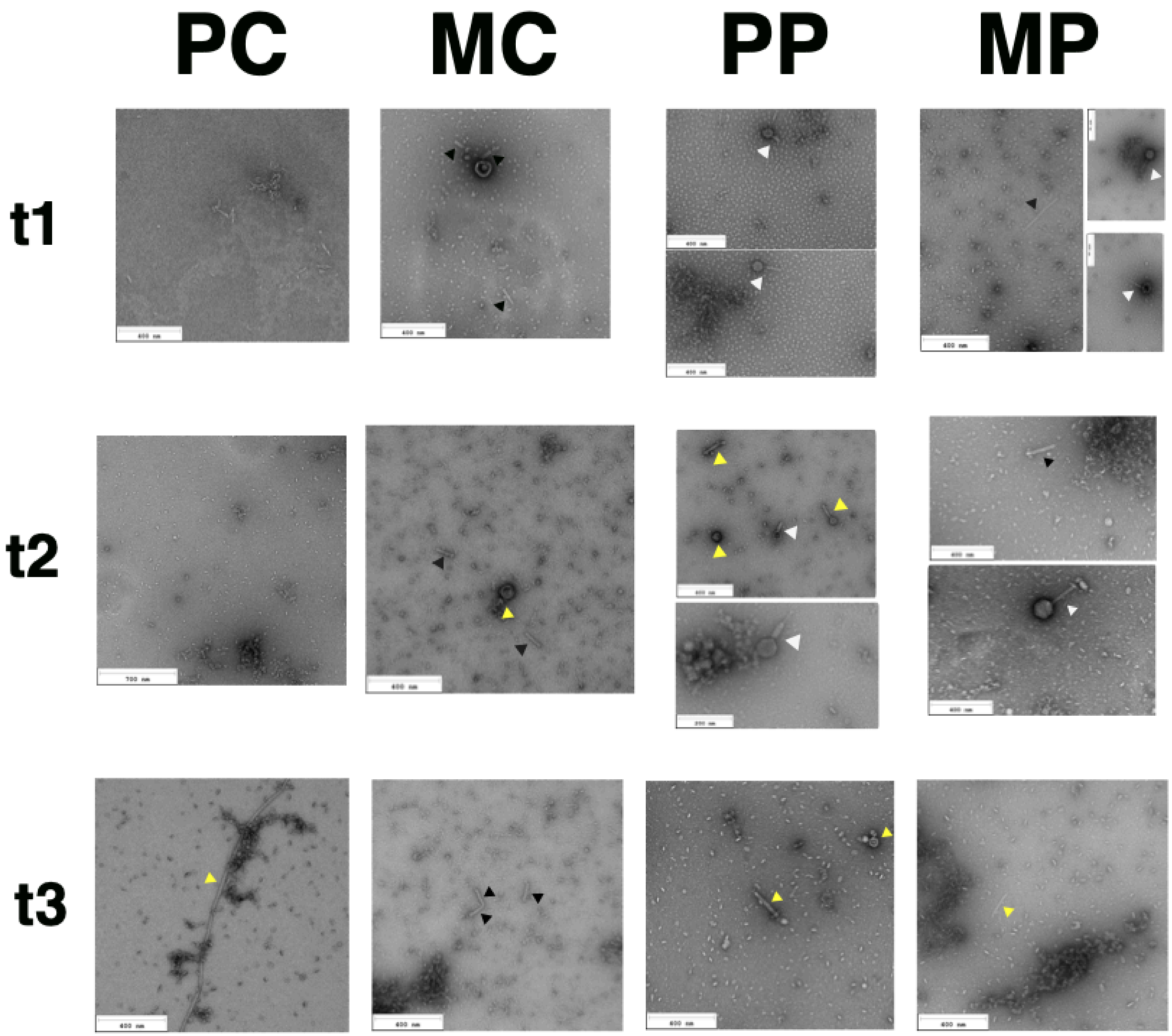


**Figure S8. Transmission electron microscopy images of *in vitro* communities filtrates.** *In vitro* communities at time points t1-t3 filtrates were used for TEM (replicate 5). White arrows denote bacteriophages; black arrows, tailocins and yellow arrows are unclear findings. MC-Mixed community, MP- Mixed community with phages, PC- Pathogenic community, PP- Pathogenic community with phages. Uncropped and additional photos for the TEM session can be found in the raw data repository.

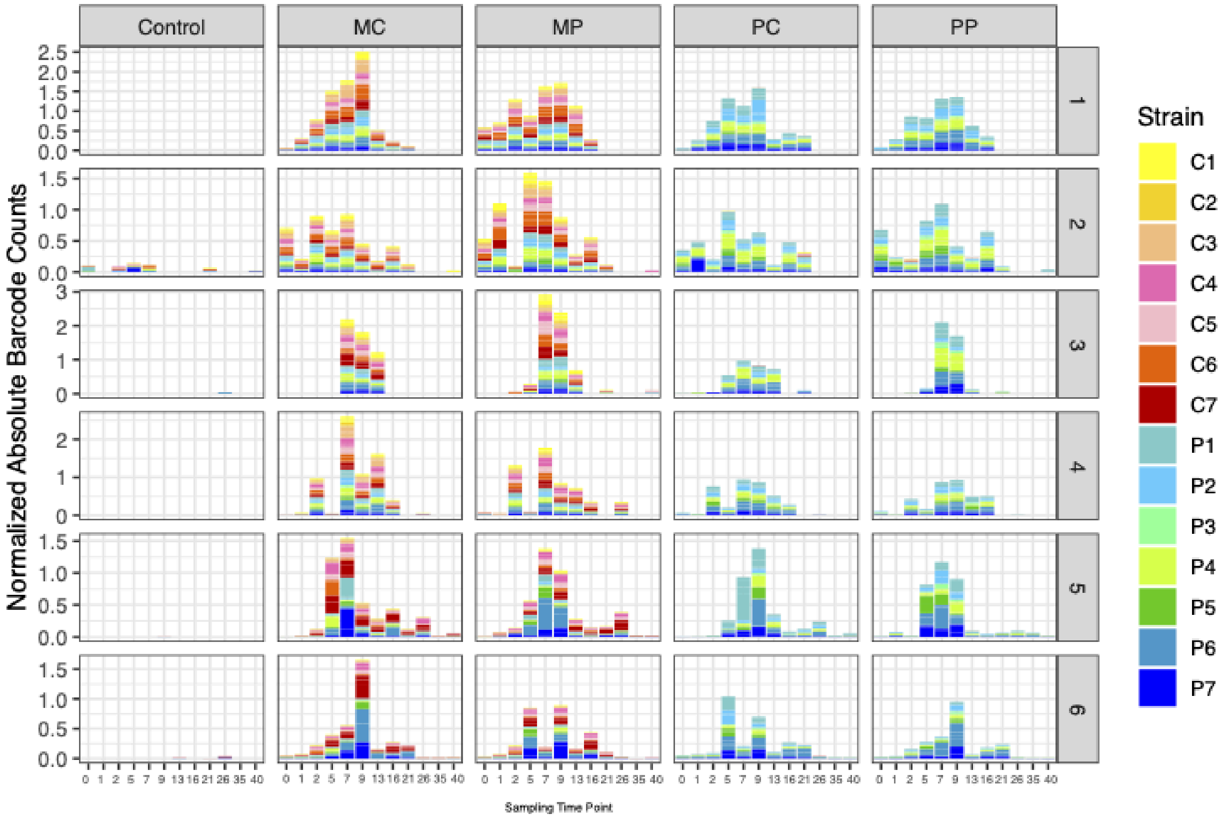


**Figure S9. Absolute abundances of bacterial strains *in planta*.** Relative abundance of each bacterial strain was calculated by amplicon sequencing of the barcode; total bacterial abundance was calculated by qPCR on the common barcode sequence (primers 79.80). The absolute abundance for each strain is the result of multiplying the specific strain relative abundance by the total abundance of barcodes in the sample. MC-Mixed community, MP- Mixed community with phages, PC- Pathogenic community, PP- Pathogenic community with phages.

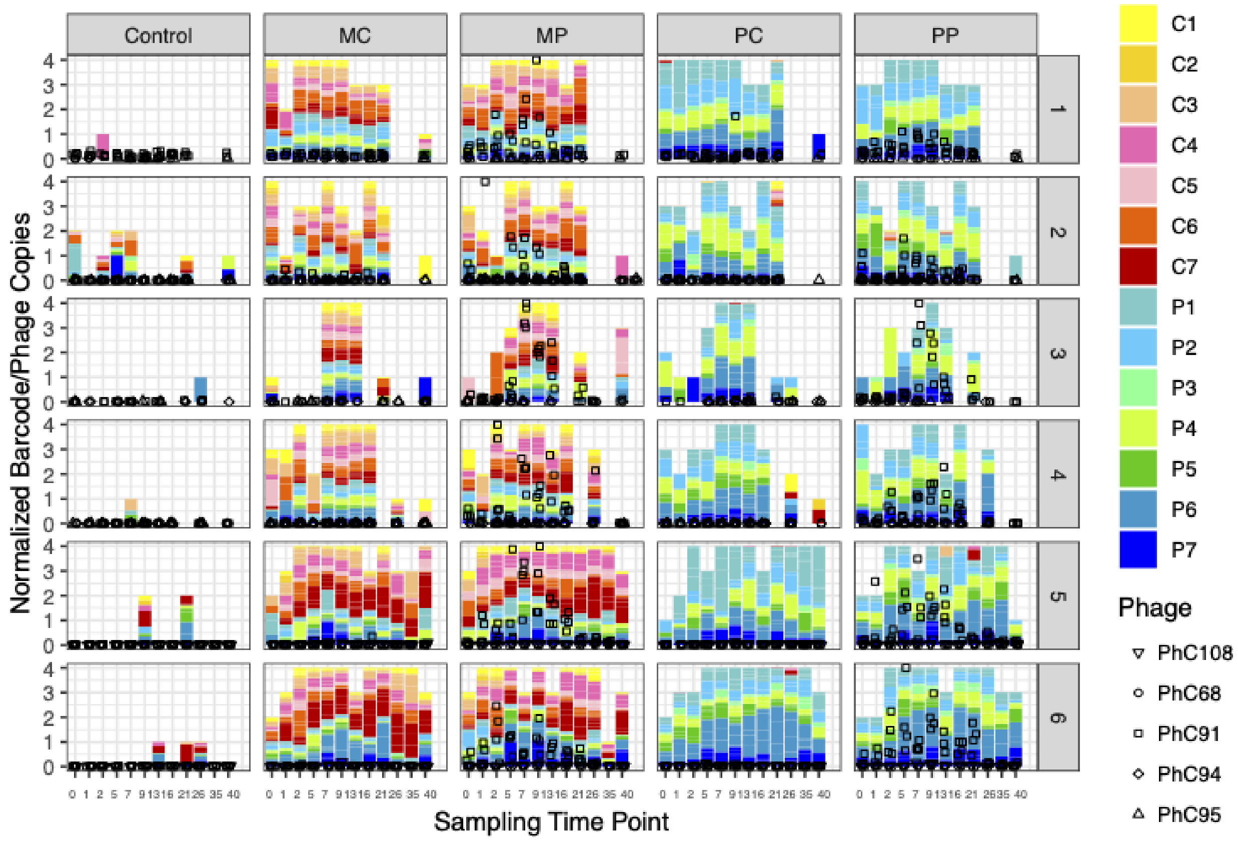


**Figure S10. Bacterial strains and phages *in planta*.** Relative abundance of the barcoded *Pseudomonas in planta*. Four plants were sampled at each time point and used for amplicon sequencing. Missing plants failed to yield reads after two sequencing runs. Symbols represent bacteriophage abundance calculated by qPCR and normalized to 4 plants per experiment. Control plants are shown for comparison with treatments, the total number of reads in control treatments were very low (see Figure S9 for absolute copies and Table S19 for non-normalized values).

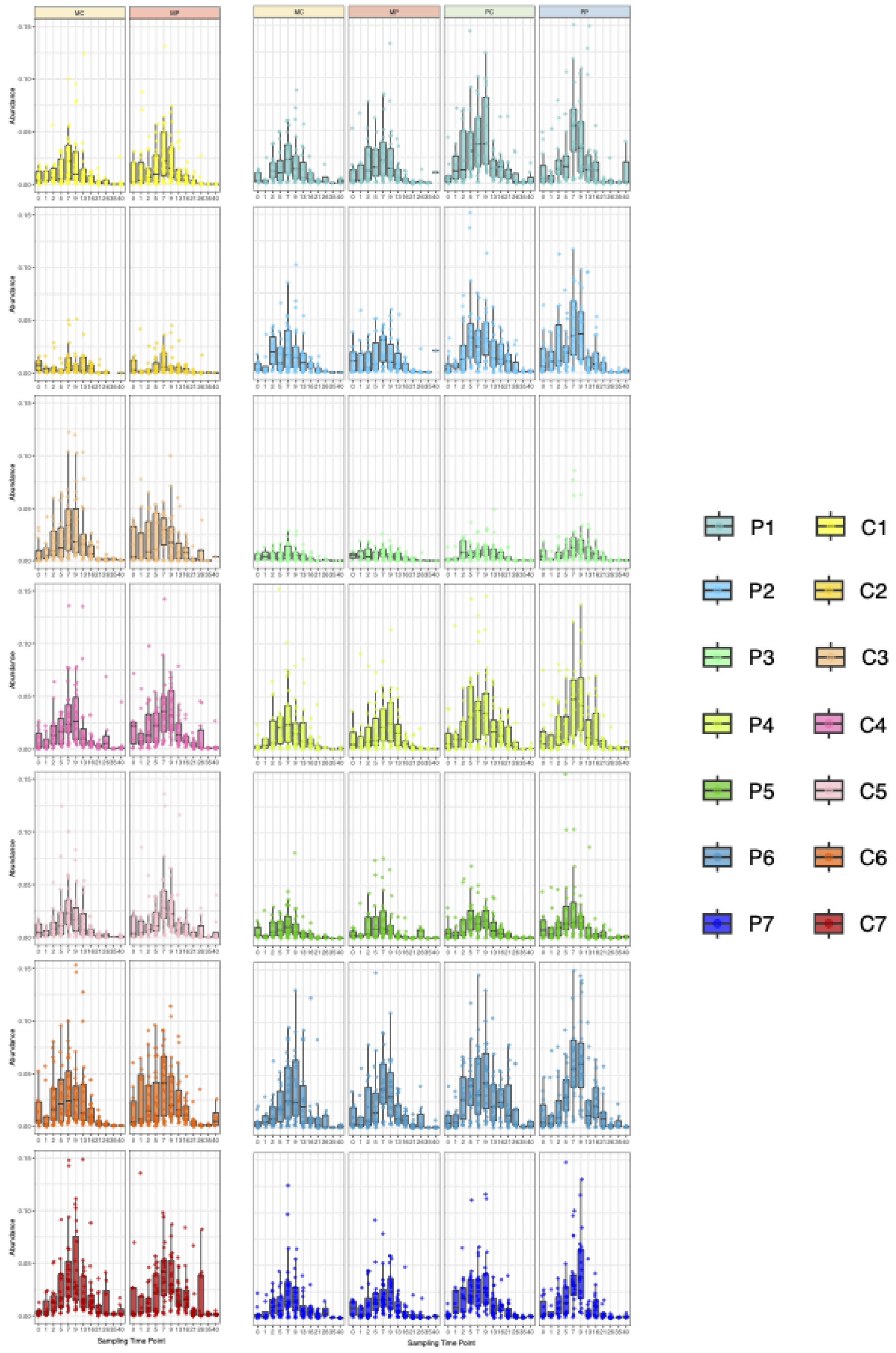


**Figure S11. *Pseudomonas* strains abundance *in planta*.** Barcoded *Pseudomonas* strains abundance per treatment. Absolute abundance was calculated by multiplying the relative abundance of the strain by the barcode copies in each sample. Y-axis maximum was uniformly set at 0.15 to accommodate all strains. MC-Mixed community, MP- Mixed community with phages, PC- Pathogenic community, PP- Pathogenic community with phages.

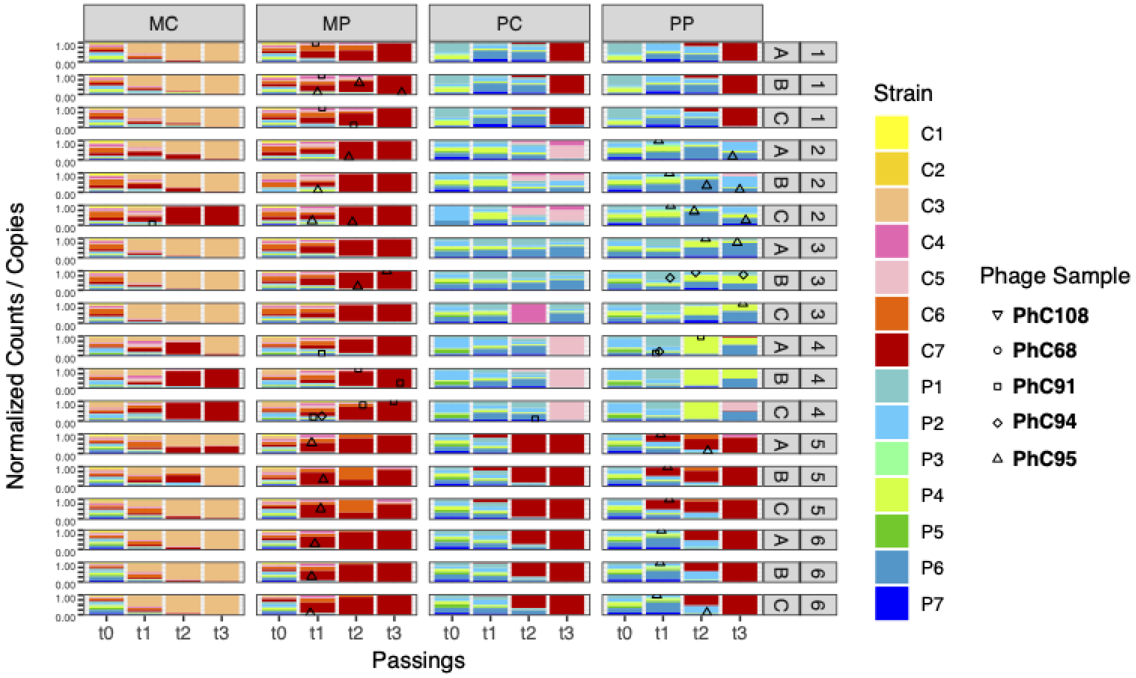


**Figure S12. Relative abundance of bacterial strains and phages in *in vitro* communities.** Relative abundance of Pseudomonas strains was calculated by amplicon sequencing; phage abundance by qPCR. Shapes and color according to the legend. X-axis denotes daily passings of the culture on a 1:10 dilution. 1-6 denote the experimental replicate, A,B,C are three biological replicates for the same experiment. MC-Mixed community, MP- Mixed community with phages, PC- Pathogenic community, PP- Pathogenic community with phages. For clarity, phage shapes are shown only for points above noise level. Data can be found in Tables S14, S15.

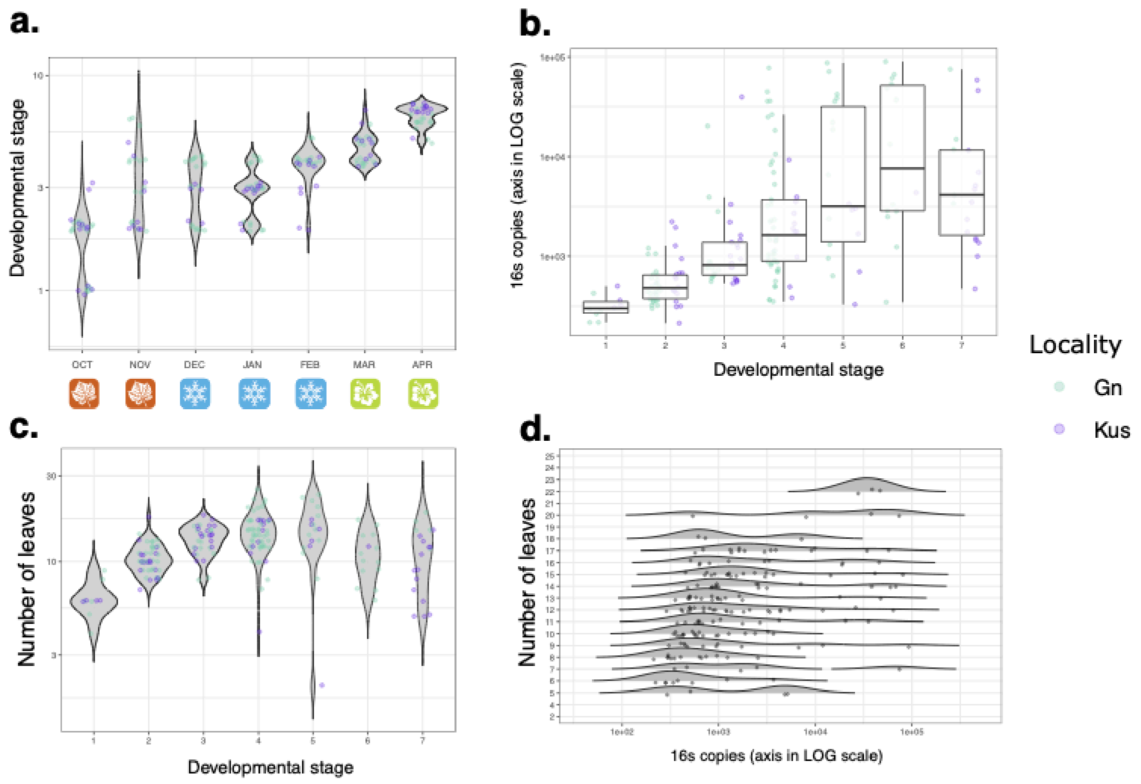


**Figure S13.** ***A. thaliana*** **wild plants traits. a.** Developmental stage of *A. thaliana* plants over the growing season. A numerical value for developmental stage was given to each plant, ranging from Juvenile (1) to flowering (7). **b.** Bacterial abundance (16S rDNA copies calculated by qPCR) on plants per developmental stage, colored by population. Y-axis is presented in log scale to facilitate visualization. **c.** Number of leaves in a plant per developmental stage. **d.** Bacterial abundance per plant categorized by its leaves number. X-axis is presented in log scale to facilitate visualization.

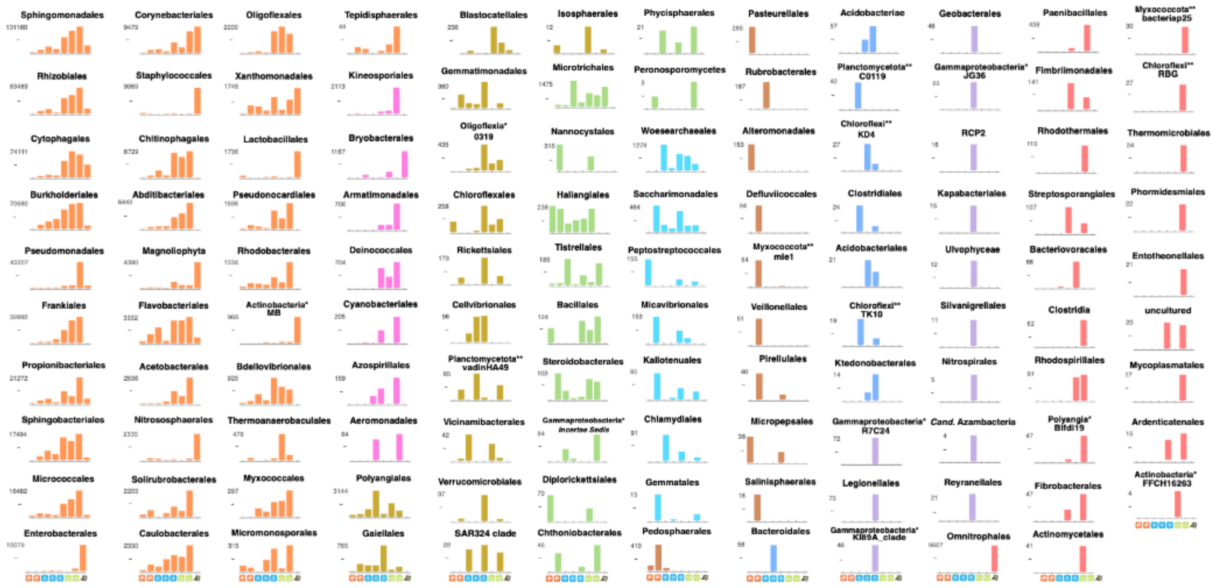


**Figure S14. Bacterial orders diversity and abundance in wild *A. thaliana* populations over the growing season.** Bacterial diversity was measured by 16S rDNA amplicon sequencing for each plant, and their relative abundance multiplied by the corresponding value of 16S rDNA total copies, to obtain the total abundance of each family. For easier visualization the figure was created at the order taxonomic level, counts can be found in Table S20. Orders are grouped and colored by similar dynamics over the season: Orange – orders present in most sampling times, increasing in abundance over the season; magenta – partially present, increase in abundance over the season; gold- peaks in winter and decreases towards the end of the season; green- high abundances in autumn and spring, less abundant during winter; cyan- peak in autumn and decline. Brown columns represent orders found solely in autumn; blue denote orders detected in winter; lilac represent orders detected at the transition to spring, by the end of February; red are orders found solely in spring. Y-axis value marks the maximum column value (copies) for the graph, "-" marks the middle value. X-axis is marked by the seasons, corresponding to the end of October - April on a monthly basis, and a flower to mark flower-only samples.

**a.**

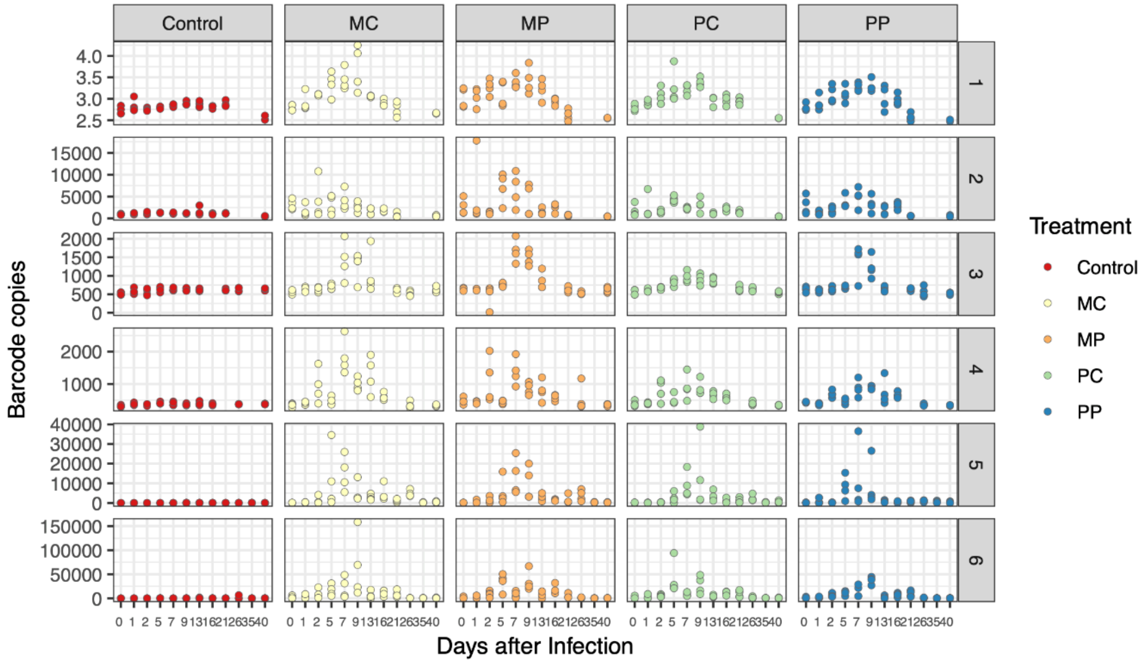


**b.**

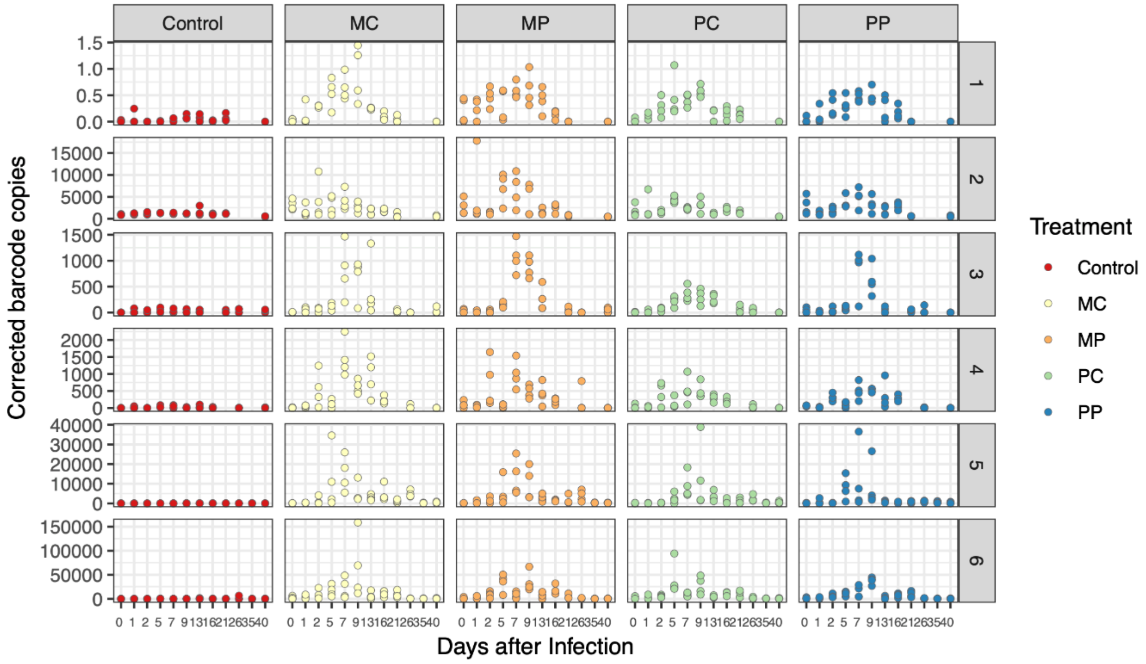


**Figure S15. Bacterial barcode abundances *in planta*.** Barcodes measured for bacterial abundances *in planta* **a.** before and **b.** after correction, based on the control treatment background. Absolute values can be found in Table S18.
